# Dynamic control over feedback regulation identifies pyruvate-ferredoxin oxidoreductase as a central metabolic enzyme in stationary phase *E. coli*

**DOI:** 10.1101/2020.07.26.219949

**Authors:** Shuai Li, Zhixia Ye, Juliana Lebeau, Eirik A. Moreb, Michael D. Lynch

## Abstract

We demonstrate the use of two-stage dynamic metabolic control to manipulate feedback regulation in central metabolism and improve stationary phase biosynthesis in engineered *E. coli*. Specifically, we report the impact of dynamic control over two enzymes: citrate synthase, and glucose-6-phosphate dehydrogenase, on stationary phase fluxes. Firstly, reduced citrate synthase levels lead to a reduction in ***α***-ketoglutarate, which is an inhibitor of sugar transport, resulting in increased stationary phase glucose uptake and glycolytic fluxes. Reduced glucose-6-phosphate dehydrogenase activity activates the SoxRS regulon and expression of pyruvate-ferredoxin oxidoreductase, which is in turn responsible for large increases in acetyl-CoA production. The combined reduction in citrate synthase and glucose-6-phosphate dehydrogenase, leads to greatly enhanced stationary phase metabolism and the improved production of citramalic acid enabling titers of 126±7g/L. These results identify pyruvate oxidation via the pyruvate-ferredoxin oxidoreductase as a “central” metabolic pathway in stationary phase *E. coli*, which coupled with ferredoxin reductase comprise a pathway whose physiologic role is maintaining NADPH levels.

**Highlights:**

- Dynamic reduction in ***α***-keto-glutarate pools alleviate inhibition of PTS dependent transport improving stationary phase sugar uptake.
- Dynamic reduction in glucose-6-phosphate dehydrogenase activates pyruvate flavodoxin/ferredoxin oxidoreductase and improves stationary acetyl-CoA flux.
- Pyruvate flavodoxin/ferredoxin oxidoreductase is responsible for large stationary phase acetyl-CoA fluxes under aerobic conditions.
- Production of citramalate to titers 126 ± 7g/L at > 90 % of theoretical yield.

## Introduction

Most metabolic engineering strategies aimed at improving strain performance rely on the overexpression of desired pathway enzymes and deletion and/or downregulation of competing biochemical activities. Over the last several decades, stoichiometric models of metabolism have helped to move the field from manipulating gene expression levels to manipulating networks, which can now be designed to couple growth with product formation, and selection can be used to optimize for both.^1,2^

A remaining limitation to these approaches are the metabolic boundary conditions required for cellular growth. Dynamic metabolic control and specifically two-stage control offer a potential engineering strategy to overcome these limitations, by switching to a production state where metabolite and enzyme levels can be pushed past the boundaries required for growth.^3–9^ Significant efforts have been made to develop tools for dynamic metabolic control including control systems, metabolic valves and modeling approaches.^3–12^ However to date, previous work has largely focused on dynamically redirecting fluxes by switching “OFF” pathways that stoichiometrically compete with a desired pathway.^6^ In this work, we demonstrate increased stationary phase flux attributable to dynamic reduction in metabolites which act as feedback regulators of central metabolism, and not reductions in competing metabolic pathways.

As illustrated in Figure 1, we investigate the impact of dynamic control of two central metabolic pathways (the tricarboxylic acid (TCA) cycle and pentose phosphate pathway (PPP)) on flux through glycolysis and pyruvate oxidation. We accomplish this by creating synthetic metabolic valves (SMVs),^3–5,13,14^ to dynamically reduce levels of the first committed step in each pathway, namely citrate synthase (GltA, “G”, encoded by the *gltA* gene) and glucose 6-phosphate dehydrogenase (Zwf, “Z”, encoded by the *zwf* gene). We show that the dynamic control over these two enzymes improves stationary phase production of citramalate, and has applicability in the production of numerous products requiring pyruvate and/or acetyl-CoA. Importantly, we also demonstrate that these improvements in flux are attributable to pyruvate ferredoxin oxidoreductase (Pfo), highlighting the importance of this enzyme in central metabolism in the stationary phase. The magnitude of flux through Pfo, an oxygen labile enzyme often thought to be active only in anaerobic conditions, is both unexpected and an example of the uncharacterized metabolic potential of even the most well characterized microbes.

**Figure 1:**
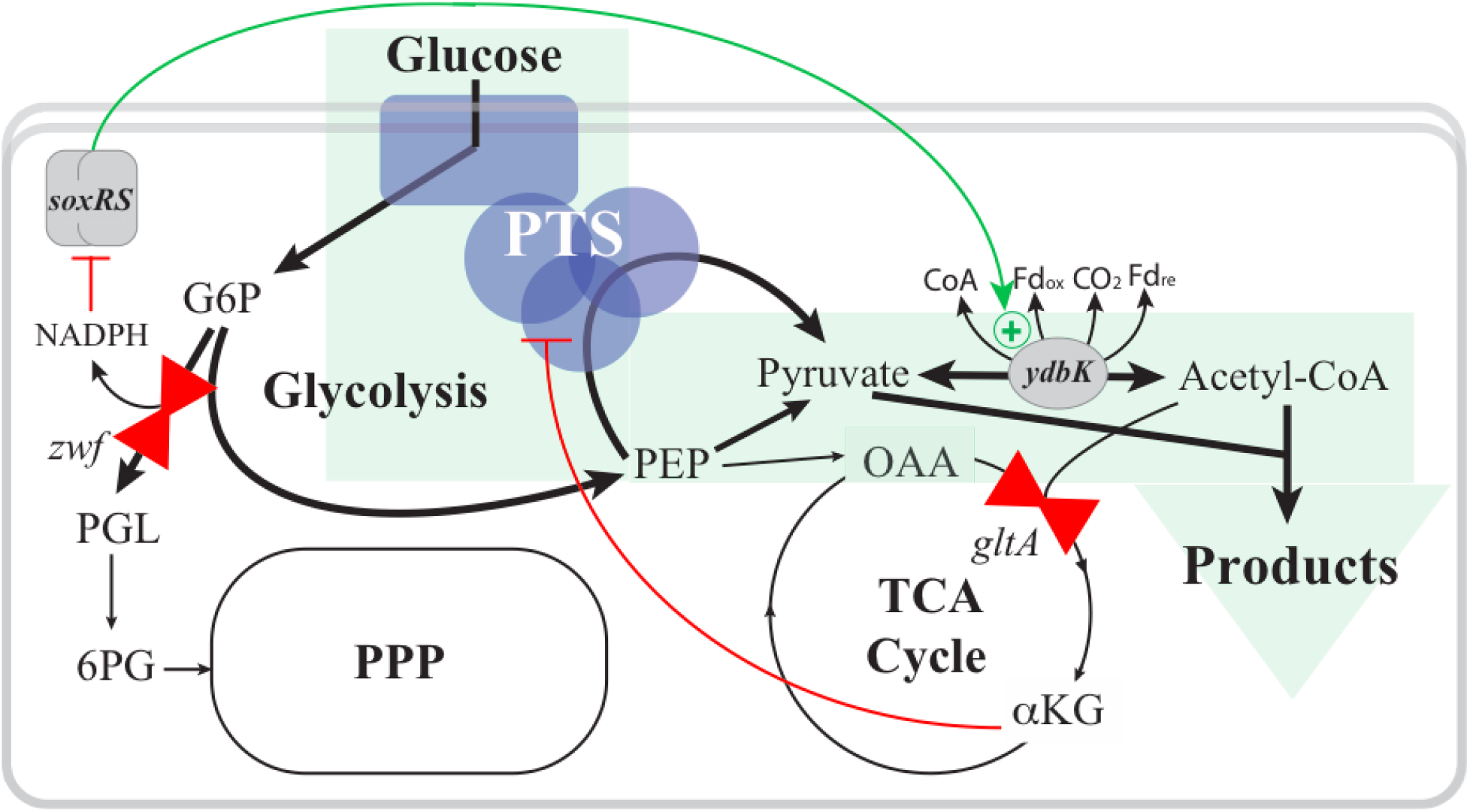
Two-stage dynamic control over feedback regulation of central metabolism improves stationary phase sugar uptake and acetyl-CoA flux. Metabolic valves (red double triangles) dynamically reduce levels off Zwf (glucose-6-phosphate dehydrogenase) and GltA (citrate synthase). Reduced flux through the TCA cycle reduces *α*KG levels alleviating feedback inhibition of PTS dependent glucose uptake, improving glycolytic fluxes and pyruvate production. Reduced flux Zwf reduces NADPH levels activating the SoxRS oxidative stress response regulation and increasing expression and activity of pyruvate ferredoxin oxidoreductase improving pyruvate oxidation and acetyl-CoA flux. Abbreviations: PTS: phosphotransferase transport system, PPP: pentose phosphate pathway, TCA: tricarboxylic acid, G6P: glucose-6-phosphate, 6-PGL: 6-phosphogluconolactone, 6PG: 6-phosphogluconate, PEP: phosphoenolpyruvate, Fd: ferredoxin, CoA: coenzyme A, OAA: oxaloacetate, *α*KG: *α*-ketoglutarate.

## Results

We first developed control systems capable of the dynamic reduction of protein levels in two-stage processes, as illustrated in Figure 2. Valves rely on controlled proteolysis or CRISPRi/Cascade based gene silencing or both proteolysis and silencing in combination to reduce levels of key metabolic enzymes.^3,4,15–18^ Induction is implemented using phosphate depletion as an environmental trigger.^19–22^ The native *E. coli* Type I-E Cascade/CRISPR system is used for gene silencing (Figure 2b).^16,23^ Targeted proteolysis is implemented by linking the expression of the chaperone SspB to phosphate deprivation. SspB, when induced, binds to C-terminal DAS+4 peptide tags on any target protein and causes degradation by the ClpXP protease of *E. coli* (Figure 2c).^17^ Refer to Supplemental Tables for strains and plasmids used in this study. Using engineered strains, as Figure 2d demonstrates, protein levels can be controlled in a two-stage process, as exemplified by turning “ON” GFP and “OFF” constitutively expressed^24^ mCherry. While, in this case, the combination of gene silencing with proteolysis results in the largest rates of protein degradation (Figure 2e and f), the impact of each approach and specific decay rates, will vary depending on the target gene/enzyme and its specific natural turnover rates and expression levels.^25,26^

**Figure 2:**
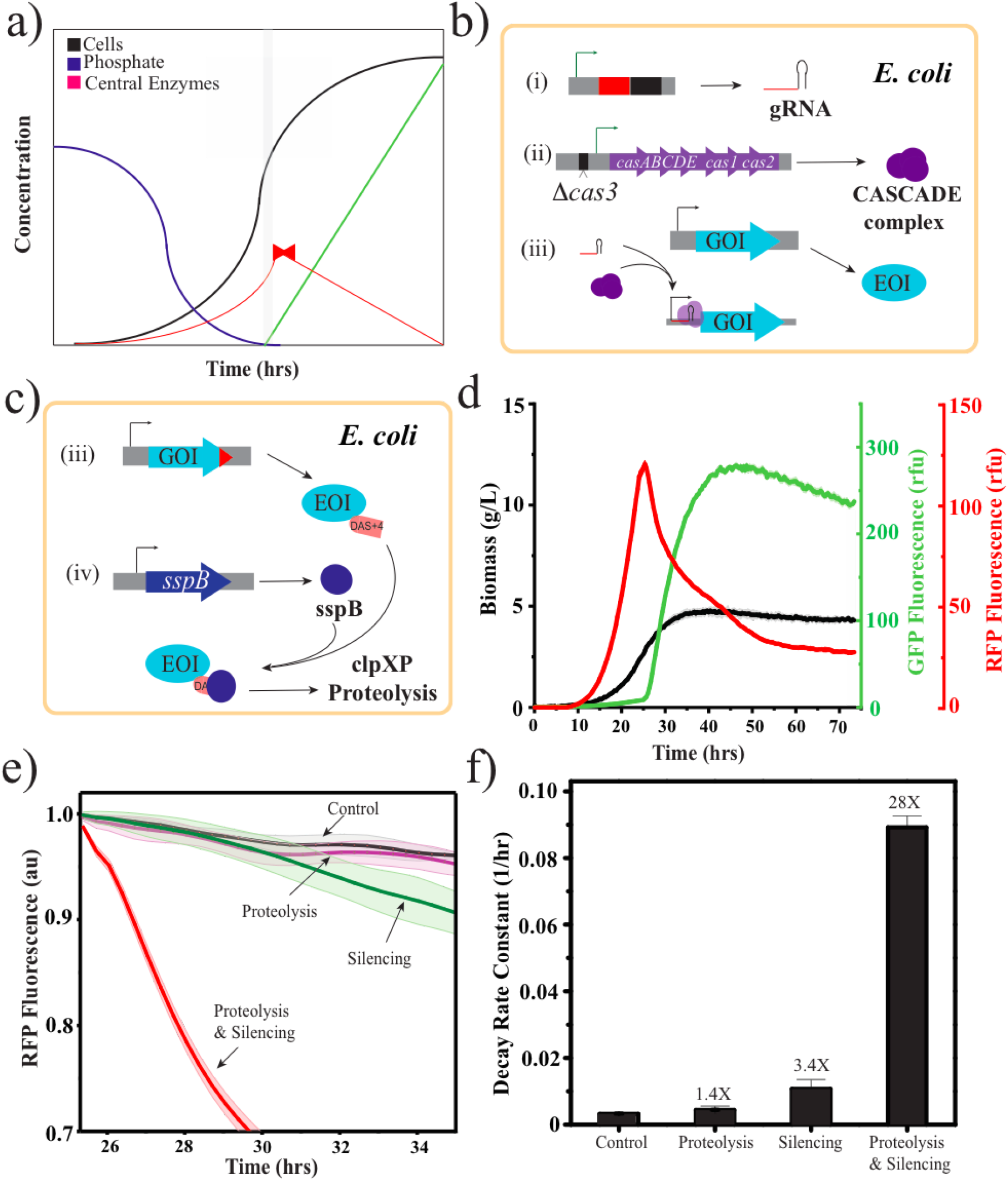
(a) Time course of two stage dynamic metabolic control upon phosphate depletion Biomass levels accumulate and consume a limiting nutrient (in this case inorganic phosphate), which when depleted triggers entry into a productive stationary phase, levels of key enzymes are dynamically reduced with synthetic metabolic valves (red) (b & c). Synthetic metabolic valves utilizing CRISPRi based gene silencing and/or controlled proteolysis. (b) Array of silencing guides can be used to silencing target multiple genes of interest (GOI). This involves the inducible expression of one or many guide RNAs as well as expression of the modified native Cascade system wherein the cas3 nuclease is deleted. The gRNA/Cascade complex binds to target sequences in the promoter region and silences transcription. (c) C-terminal DAS+4 tags are added to enzymes of interest (EOI) through chromosomal modification, they can be inducibly degraded by the clpXP protease in the presence of an inducible sspB chaperone. (d) Dynamic control over protein levels in *E. coli* using inducible proteolysis and CRISPRi silencing. As cells grow phosphate is depleted, cells “turn off’ mCherry and “turn on” GFPuv. Shaded areas represent one standard deviation from the mean, n=3. (e) The relative impact of proteolysis and gene silencing alone and in combination on mCherry degradation, (f) mCherry decays rates.

We then turned to dynamically reduce levels of GltA and Zwf (Figure 3). We engineered strains with chromosomal modifications that appended C-terminal DAS+4 degron tags to these genes. In addition, we engineered several strains to have C-terminal superfolder GFP tags behind each gene with and without C-terminal degron tags.^17,27^ Plasmids expressing gRNAs were designed to repress expression from the *gltAp2* and *zwf* promoters. Using these strains and plasmids, dynamic control over enzyme levels were monitored by tracking GFP via an ELISA assay in two-stage minimal media micro-fermentations as reported by Moreb et al.^28^ An ELISA was used as protein levels were too low in engineered strains to use GFP fluorescence as a direct reporter. In the case of GltA proteolysis and silencing resulted in a 70% and 85% decrease in GltA levels, respectively, with the combination resulting in a 90% reduction. In the case of Zwf, proteolysis, silencing as well as the combination all resulted in protein levels below the limit of quantification of our assay.

**Figure 3:**
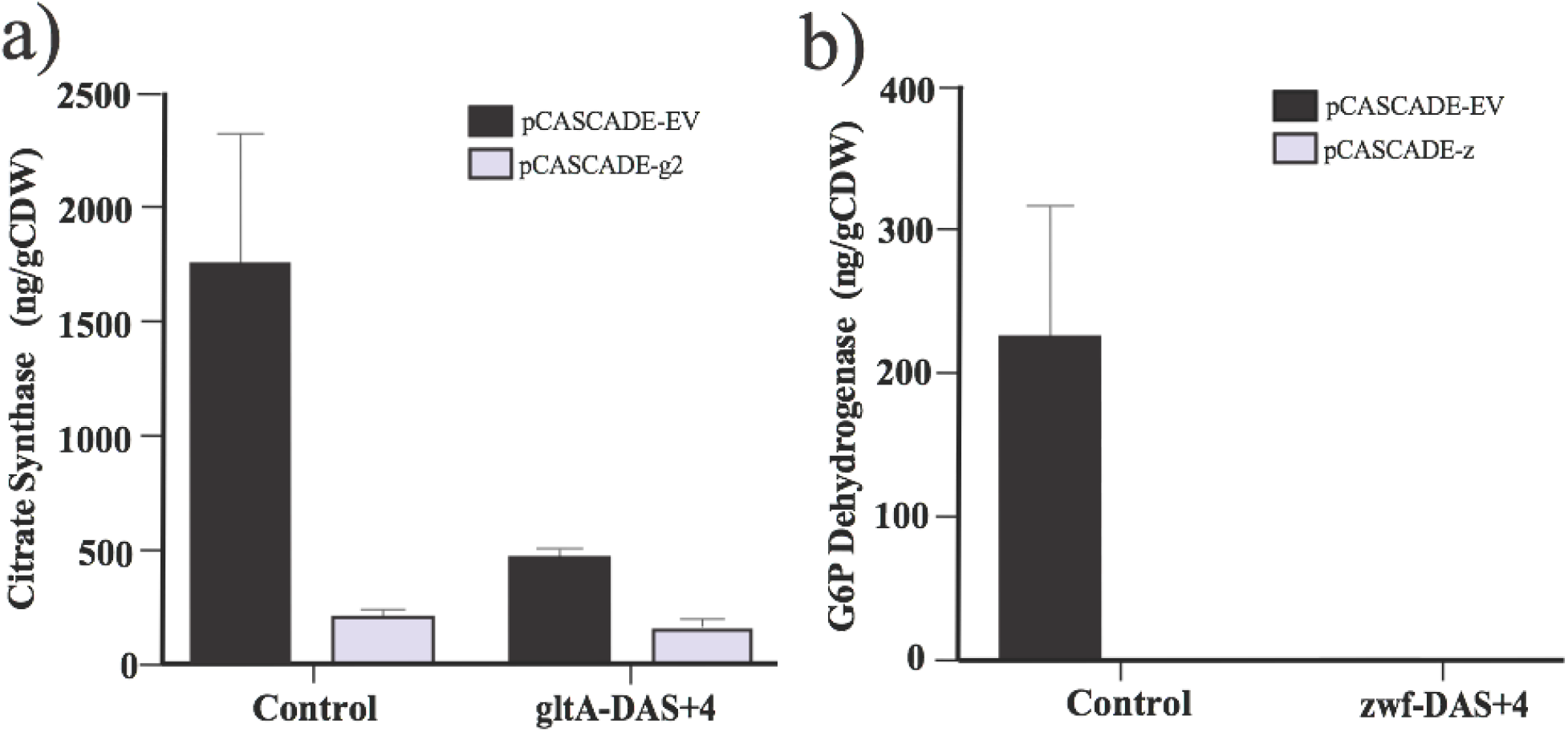
Dynamic Control over the levels of the central metabolic enzymes. The impact of silencing (pCASCADE) and proteolysis (DAS+4 tags) on protein levels were evaluated alone and in combination (a) GltA (citrate synthase), and (b) Zwf (glucose-6-phosphate dehydrogenase). In all cases chromosomal genes were tagged with a C-terminal sfGFP. Protein levels were measured by ELISA, 24 hour post induction by phosphate depletion in microfermentations.

The impact of “G” and “Z” valve combinations on metabolic fluxes were measured in minimal media micro-fermentations,^28^ performed without any heterologous production pathway. As the strains used had deletions in the major pathways leading to acetate production (*poxB*, and *pta-ackA*), pyruvate synthesis was initially evaluated as a measure of metabolic fluxes through glycolysis (Figure 4). The “G” valve had the largest impact on pyruvate production, with no detectable product measured in a control strain without SMVs. The improved production of pyruvate could be attributable either to a stoichiometric effect, wherein a portion of flux normally entering the TCA cycle is redistributed to the overflow metabolite, or alternatively to a more global increase in the sugar uptake rate enabling greater overflow metabolism and pyruvate synthesis. To evaluate these two alternatives we measured the impact of the “G” valve on glucose uptake rates. Results, shown in Figure 4c, indicate that increases in pyruvate production are primarily attributed to increases in uptake rates rather than a repartitioning of basal fluxes.

**Figure 4:**
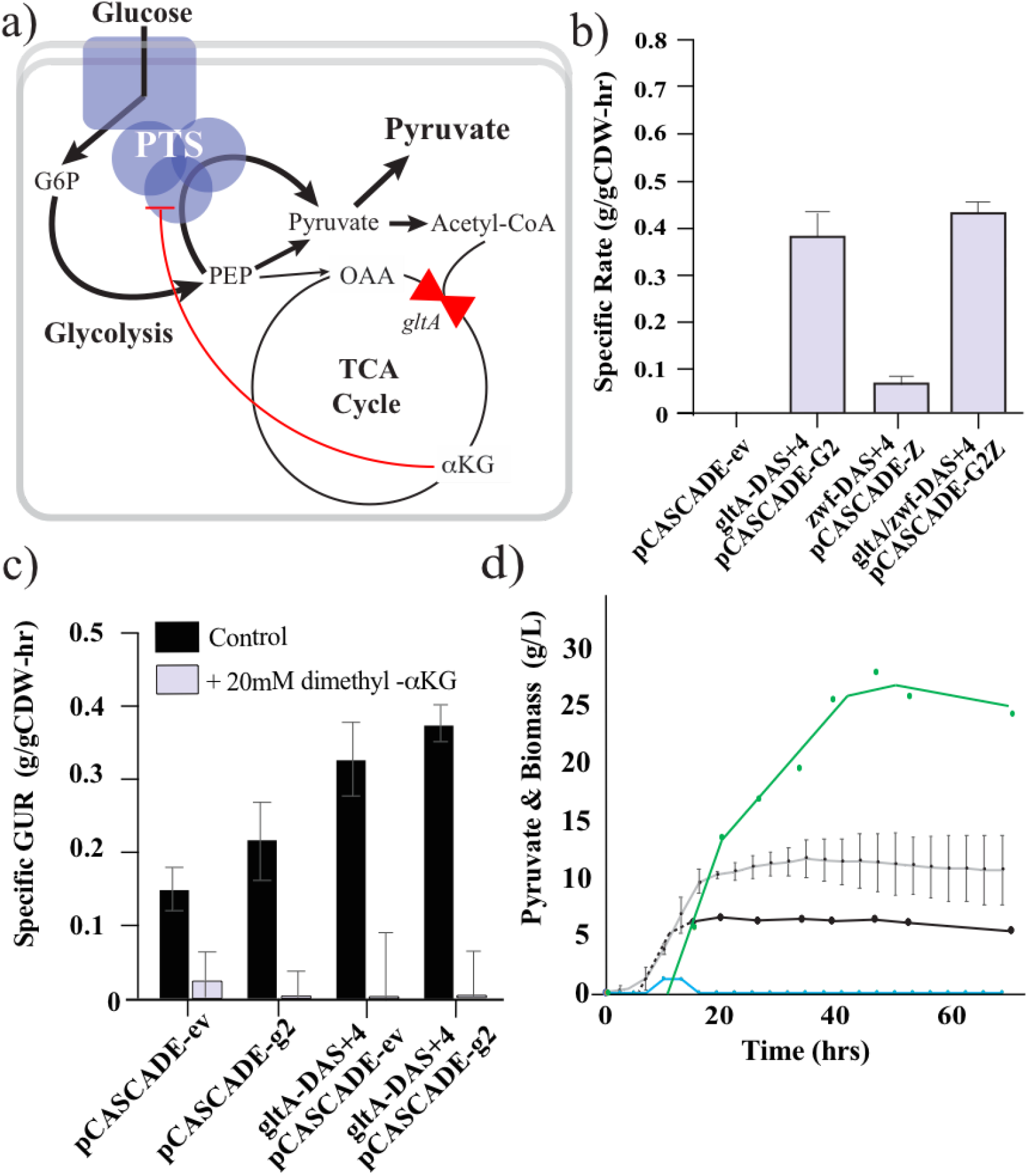
a) Dynamic reduction in GltA reduces *α*KG pools and alleviates *α*KG mediated inhibition of PTS-dependent glucose uptake (specifically, PtsI), improving glucose uptake rates, glycolytic fluxes and pyruvate production. b) The impact of dynamic control over GltA and Zwf levels on pyruvate production in minimal media microfermetnations. c) The impact of dynamic control over GltA and Zwf levels and dimethyl-*α*KG supplementation on glucose uptake rates in microfermentations. (d) Pyruvate and biomass production were measured for the control strain and the “G” valve strain. The control strain’s biomass (gray) and pyruvate production (blue), as well as the “G” valve strain’s biomass (black) and pyruvate production (green) are plotted as a function of time. Dashed line represents extrapolated growth due to missed samples.

We hypothesized increased sugar uptake due to the “G” valves was due to a direct regulatory effect of metabolites produced by the TCA cycle, namely α-ketoglutarate (αKG). αKG, a precursor to glutamic acid, has several key regulatory roles,^29–31^ including the regulation of sugar transport by direct inhibition of Enzyme I of the PTS dependent glucose transporter (Figure 1). This feedback regulation is a way to coordinate sugar uptake with nitrogen assimilation (glutamate synthesis).^31^ To test our hypothesis, we performed supplementation experiments, spiking 20mM dimethyl-αKG (DM-αKG) into microfermentations at the onset of production.^31^ DM-αKG, rather than αKG, was used as it has been shown to better cross the membrane, and then be hydrolyzed to αKG.^31^ As seen in Figure 4c, DM-αKG inhibited sugar uptake in control cells as well as in strains with valves reducing GltA levels. Together these results support dynamic reduction in GltA levels and the subsequent reduction in αKG pools as primarily responsible for improved sugar uptake rates and pyruvate biosynthesis. We next turned to assess pyruvate production in instrumented bioreactors. Minimal media fed batch fermentations were performed as previously reported by Menacho-Melgar et al.,^32^ where phosphate concentration limited biomass levels and once was consumed expression of the silencing gRNAs and the SspB chaperone are induced. Results comparing the control host strain with a strain having dynamic control over GltA levels are given in Figure 4d. Minimal pyruvate transiently accumulated in the control strain whereas maximal titers of over ~ 30g/L were obtained using dynamic control.

To assess the impact of dynamic control over acetyl-CoA fluxes we leveraged citramalate synthase which produces one mole of citramalate from one mole of pyruvate and one mole of acetyl-CoA (Figure 5a).^32,33^ Citramalate is a precursor to the industrial chemicals itaconic acid and methyl methacrylate, as well as an intermediate in branched chain amino acid biosynthesis.^31–39^ To produce citramalate, we used a low phosphate inducible plasmid expressing a previously reported feedback resistant mutant citramalate synthase (cimA3.7).^32,38,40^ This plasmid was introduced into the set of “G” and “Z” valve strains which were then assessed for citramalate production in two stage micro-fermentations (Figure 5b). The best producing strain had both “G” and “Z” valves. Next, we evaluated citramalate production strains in instrumented bioreactors. The control strain made reasonable citramalate titers (~40g/L), whereas the introduction of SMVs improved production. The combined “GZ” valve strain had the highest citramalate production, reaching titers of ~100 g/L. This process was then intensified, by increasing biomass levels from ~ 10gCDW/L to ~ 25gCDW/L, leading to titers of 126+/-7g/L. This process is illustrated in Figure 5d. Refer to Supplemental Figure S9 for the time courses for additional fermentations. The overall process yields were 0.74-0.77g citramalate/g glucose and during the production phase yields approached theoretical achieving 0.80-0.82 g citramalate/g glucose. The theoretical yield for citramalate from glucose is 1 mole/mole or 0.817g/g.

**Figure 5:**
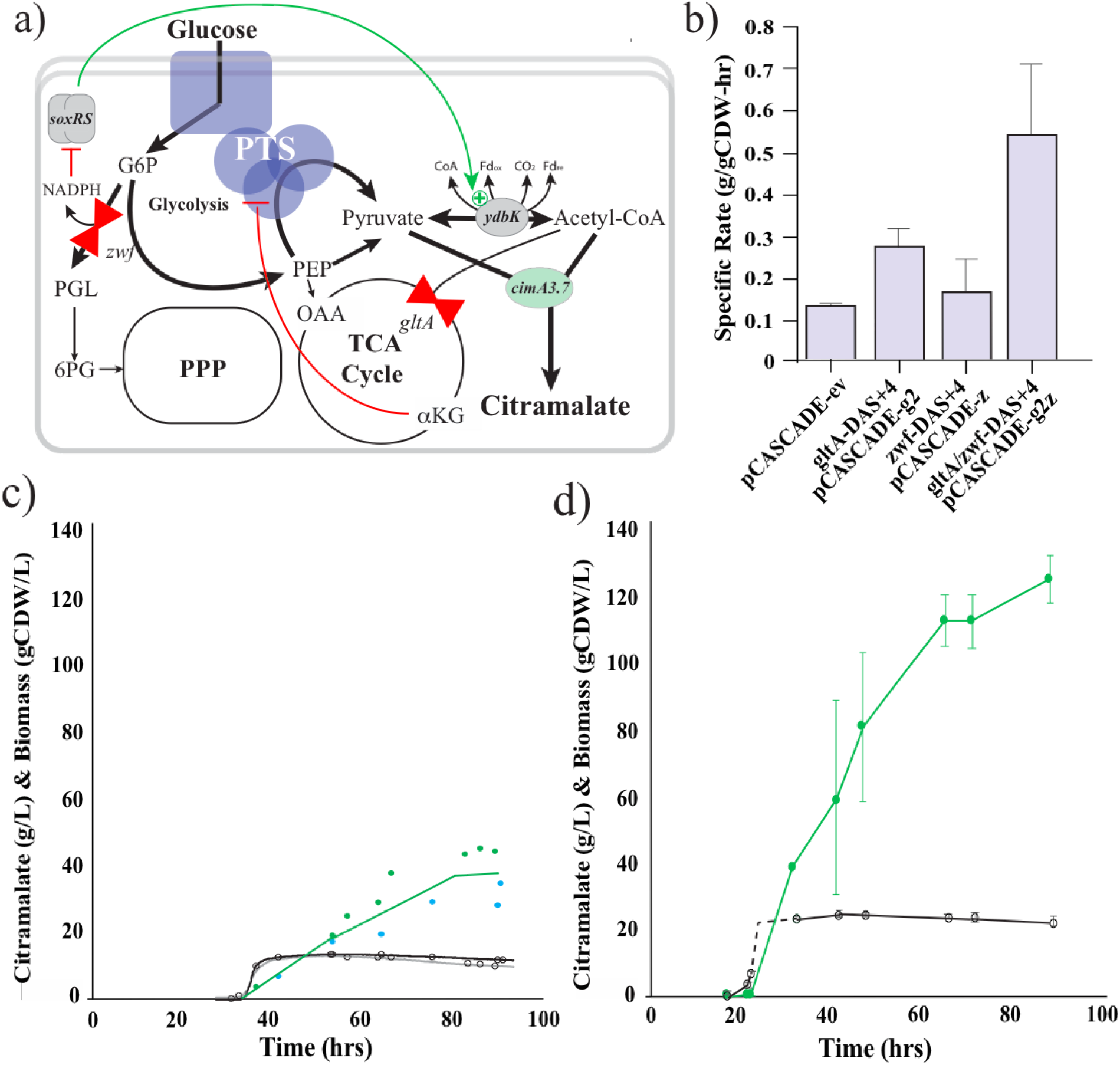
a) Dynamic reduction in Zwf levels activates the SoxRS regulon and increases activity of the pyruvate-ferredoxin oxidoreductase (Pfo, *ydbK*) improving acetyl-CoA fluxes and citramalate production. b) The impact of dynamic control over GltA and Zwf levels on citramalate production in minimal media microfermetnations. Citramalate and biomass production were measured for the control strain (c) and the “GZ” valve strain (d). (c) Duplicate runs, biomass levels in gray and black, citramalate titers in green and blue. (d) The average of triplicate runs, biomass black and citramalate green. Dashed line represents extrapolated growth due to missed samples.

In the case of pyruvate, the “Z” valve had no significant impact on production (Figure 4b). Citramalate and pyruvate are similar products in that they are both oxidized and require no redox cofactor (such as NADPH) for biosynthesis. A key difference in the two products is that citramalate requires an additional precursor, namely acetyl-CoA. We hypothesized that the “Z-valve” dependent improvement in citramalate production was dependent on improved acetyl-CoA production in strains with reduced Zwf activity. If correct, this would suggest that either Zwf levels or the levels of downstream metabolites have a negative regulatory impact on stationary phase acetyl-CoA synthesis. It is important to note that the strains used for pyruvate and citramalate production have deletions in *poxB* and *pflB* (which can lead to acetyl-CoA synthesis) and initially we had assumed all acetyl-CoA flux was through the well characterized pyruvate dehydrogenase (PDH) multienzyme complex.^41,42^ To our surprise, as shown in Figure 6a, proteolytic degradation of Lpd (a subunit of PDH) had no impact on citramalate production. Based on the this we turned to consider the potential of an alternative primary route for acetyl-CoA production in stationary phase cultures, namely pyruvate-flavodoxin/ferredoxin oxidoreductase (Pfo), encoded by the *ydbK* gene.

**Figure 6:**
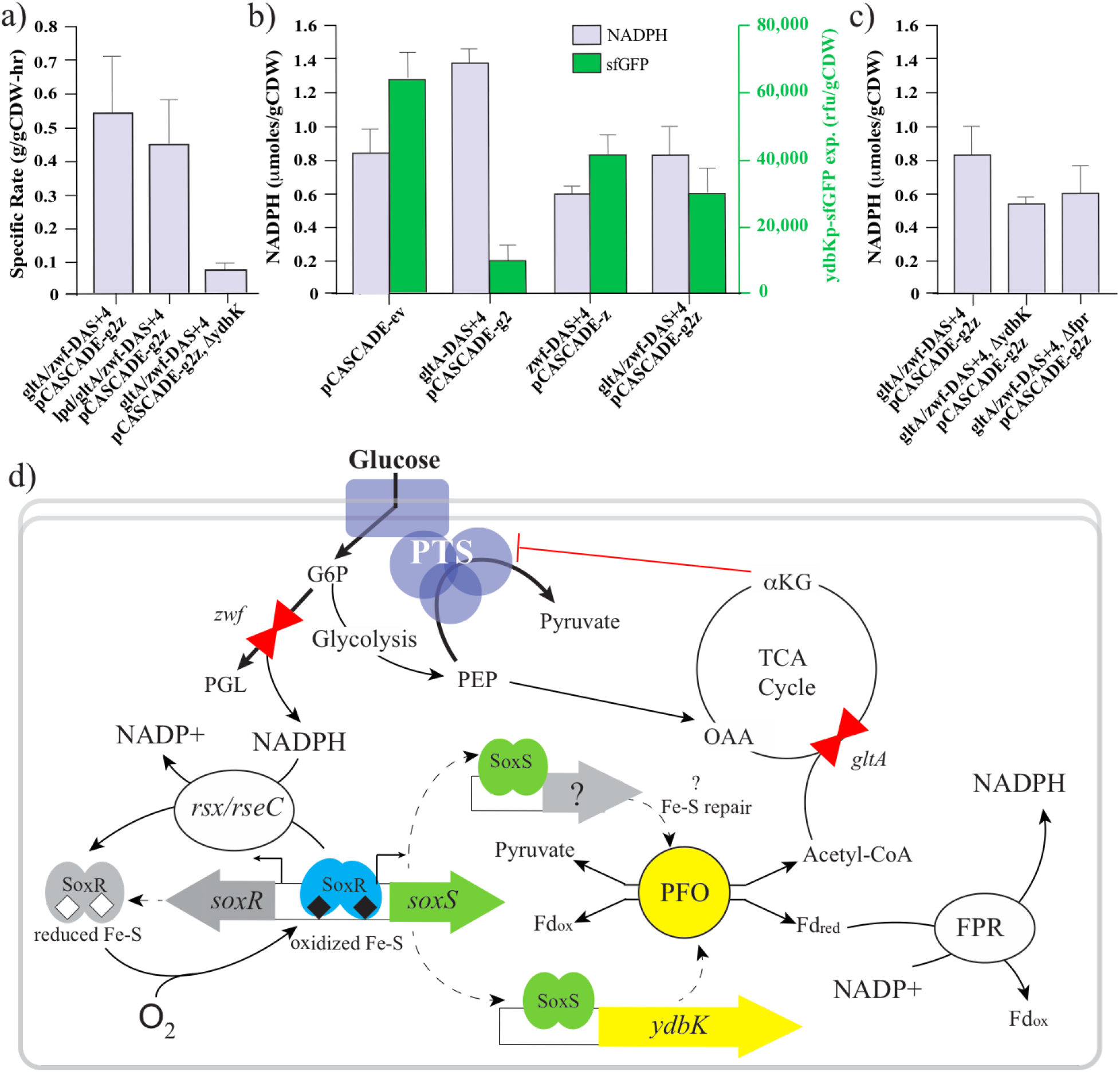
a) acetyl-CoA flux is dependent on Pfo (YdbK) activity. The proteolytic degradation of Lpd (lpd-DAS+4, a subunit of the pyruvate dehydrogenase multienzyme complex) and a deletion in *ydbK* were assessed in the “GZ” valve background. b) NADPH pools (gray bars) and ydbK expression levels (green bars) in engineered strains. Expression of a superfolder GFP (sfGFP) reporter is driven by the *ydbK* promoter. c) NADPH levels in the “GZ” valve strain as well as strains with deletions in either *ydbK* or *fpr*. d) A model for the synergistic activity of “G” and “Z” valves on acetyl-CoA production through Pfo. A combination of the “G” valve and “Z” valve are needed to i) increase sugar uptake rates and ii) reduce NADPH pools. Decreased NADPH pools lead to lower rates of reduction of the soxR iron-sulfur cluster, which is spontaneously oxidized. Oxidized soxR activated expression of *soxS*, which in turn leads to increased expression of *ydbK* and Pfo activity. We hypothesize that the oxygen labile Pfo is continually reactivated in vivo by one or more enzymes that may also be subject to soxSR regulation. Reduced ferredoxin can be used to generate NADPH *in vivo* through the action of Fpr.

We hypothesized, as illustrated in Figure 1 (as well as 6d), that Pfo (*ydbK*) was in part responsible for acetyl-CoA synthesis in stationary phase and that due to its role in the oxidative stress response this activity was regulated by intermediates in the PPP, also known to be involved in the response to oxidative stress.^43–45^ To test this hypothesis we constructed a *ydbK* deletion in the citramalate strain containing both “G” and “Z” valves and measured citramalate production. As seen in Figure 6a, the deletion of *ydbK* significantly reduced citramalate synthesis confirming the role of Pfo in acetyl-CoA flux. As Pfo has been shown to be induced upon oxidative stress, via the SoxRS regulon (which is regulated by oxidant levels and NADPH pools),^46–49^ we hypothesized that expression was due to alterations in NADPH levels caused by reductions in Zwf activity. The iron-sulfur cluster of SoxR is continually oxidized under aerobic conditions and continually reduced via at least two NADPH dependent enzymes (Rsx, RseC).^50^ These reductions are known to be sensitive to NADPH levels or pools. Previous work has shown a modest activation of SoxRS in response to a *zwf* deletion,^51^ and an impaired response to oxidative stress,^52^ which may be magnified by a rapid dynamic reduction of Zwf activity. To test this hypothesis, we performed additional confirmatory studies.

We constructed a plasmid with a superfolder GFP driven by the *ydbK* gene promoter and evaluated expression in strains with “G” and “Z” valves alone and in combination. Results are given in Figure 6b. Interestingly, the control strain had the highest expression from the *ydbK* promoter, indicating the natural activity of this enzyme in phosphate limited stationary phase. *YdbK* promoter expression was greatly reduced in the “G” valve strain. The “Z” valve strain had reasonable *ydbK* promoter expression although decreased from the control, and the combined “GZ” valve recovered significant expression when compared to the “G” valve alone. We hypothesize, as illustrated in Figure 6d, that under normal conditions, inhibition of sugar uptake generally reduces metabolism, including flux through Zwf, leading to reduced NADPH pools and activation of the SoxRS regulon. Increased sugar uptake in response to the “G” valve leads to increased flux through central metabolism (including through Zwf) which results in elevated NADPH pools and reduced SoxRS activation. The addition of the “Z” valve in combination with the “G” valve eliminates flux through Zwf reducing NADPH levels recovering SoxRS activation and *ydbK* expression. Thus this combination of “G” and “Z” valves uniquely increases sugar uptake and enables soxRS activation leading to improved acetyl-CoA flux. Stationary NADPH pools are consistent with this hypothesis. As shown in Figure 6b, the “G” valve led to increased NADPH levels compared to the control, whereas the addition of the “Z” reduced these levels, regaining *ydbK* expression.

Lastly, we wanted to further confirm the physiologic role for Pfo and its activation. We hypothesized that unlike suggestions in previous reports, Pfo has a direct role in the production of NADPH and that reduced ferredoxin/flavodoxin can be used to generate NADPH through the action of ferredoxin reductase (Fpr) operating as a dehydrogenase. The expression of*fpr* is also activated by SoxRS,^53^ although it has been thought that physiologically, Pfo and Fpr serve similar roles in generating reduced ferredoxin/flavodoxin,^54^ rather than forming a two step enzymatic pathway to produce NADPH. To test our hypothesis we measured NADPH levels in our “GZ” valve strain (eliminating or reducing NADPH production from the pentose phosphate pathway and TCA cycles) with deletions in either the *ydbK* or *fpr* genes. Results are given in Figure 6c. Either deletion resulted in decreases in NADPH levels supporting our hypothesis that Pfo and Fpr combined are capable of producing NADPH *in vivo*. If the role of Fpr was to consume NADPH to generate reduced ferredoxin/flavodoxin we would have expected a deletion to increased NADPH pools. These results are consistent with our model as illustrated in Figure 6d, wherein Pfo along with Fpr couple pyruvate oxidation and acetyl-CoA production with NADPH generation.

## Discussion

Previous studies utilizing dynamic control have primarily been informed by a stoichiometric framework, wherein pathways are switched “ON” and “OFF” to reduce fluxes that stoichiometrically compete for a desired product, or in other words pathway redirection.^6,55^ For example Venayak and colleagues^7^ have highlighted the importance of GltA/CS as a central valve candidate for dynamic metabolic control, based in part on stoichiometric modelling.^7,56^ However these studies and models have missed the importance of the regulatory role of downstream metabolites, such as αKG. This work demonstrates that increasing flux by dysregulation of feedback control can have a large impact on production, independent of stoichiometry or the minimization of competing pathways. In particular, it was unexpected that reducing Zwf activity increased acetyl-CoA fluxes.

To our knowledge this is the first report demonstrating the magnitude of the metabolic flux through Pfo and the central role of this enzyme in stationary phase metabolism. This result is somewhat unexpected. Although Pfo, an iron sulfur cluster containing enzyme, has been successfully expressed in both aerobic and anaerobic conditions,^46,48,49,57^ it is quickly inactivated by molecular oxygen *in vitro*, and as a result, conventional wisdom would suggest it is unlikely to support these types of fluxes.^46,47^ These data demonstrate that high levels of Pfo activity can be maintained even aerobically *in vivo*. As discussed above, more work is needed to understand the mechanisms underlying the observed *in vivo* oxygen tolerance of this pathway. Improved understanding may lead to alternative strategies (independent of decreasing Zwf levels or *soxS* overexpression) for optimizing flux through this pathway. Additionally, while Nakayama and colleagues proposed a role for Pfo in combination with Fpr in the oxidative stress response,^46^ this study identifies Pfo coupled with Fpr as a route for NADPH generation in *E. coli*, demonstrating that NADPH dependent ferredoxin reductase can operate physiologically as an NADP^+^-dependent reduced ferredoxin dehydrogenase, generating NADPH, which is in the opposite direction of that previously considered. This pathway is activated upon oxidative stress when NADPH generation from glucose-6-phosphate dehydrogenase is insufficient to meet cellular requirements. While more work is needed to understand the conditions enabling the reversible flux through Fpr (such as the NADPH/NADP^+^ ratio required), in effect this pathway, is a novel route to produce NADPH and acetyl-CoA from pyruvate bypassing the pyruvate dehydrogenase multienzyme complex.

More generally, this work highlights the potential of manipulating known and unknown feedback regulatory mechanisms to improve *in vivo* enzyme activities and metabolic fluxes. This approach can open numerous novel engineering strategies, and has the potential to lead to significant improvements in production rates, titers and yields. Furthermore these results confirm the metabolic potential of stationary phase cultures,^3,4,21,32^ and that there remain significant uncharacterized differences in stationary phase metabolism as compared to that of exponential growth, even in well modelled microbes such as *E. coli*. Dynamic metabolic control in two-stage cultures is uniquely suited to implement these strategies. Simply overexpressing key enzymes does not bypass native regulation and the complete removal of central metabolic enzymes and/or metabolites will often lead to growth defects and strains which need to evolve compensatory metabolic changes to meet the demands of growth. In contrast, changes to levels of central regulatory metabolites in stationary phase enable rewiring of the regulatory network and metabolic fluxes without this constraint.^3,4^

## Methods

### Media & Reagents

Unless otherwise stated, all materials and reagents were purchased from Sigma (St. Louis, MO). Luria Broth Lennox formulation was used for routine strain and plasmid propagation and construction. FGM1, FGM30, and SM10++ seed media were prepared as previously described Menacho-Melgar et al.^32^ SM10++ and SM10 no phosphate media were prepared as described by Moreb et al.^33^ FGM3 media used in biolector studies was detailed in Supplemental Materials. Working antibiotic concentrations were as follows: kanamycin: 35 μg/mL, chloramphenicol: 35 μg/mL, zeocin: 100 μg/mL, blasticidin: 100 μg/mL, spectinomycin: 25 μg/mL, tetracycline:5 μg/mL.

### Strains & Plasmids

Plasmid and strain information is given in Supplemental Materials. Sequences of oligonucleotides and synthetic linear DNA (Gblocks™) are given in Supplemental Materials, Tables S2 and S3, and they were obtained from Integrated DNA Technologies (IDT, Coralville, IA). Deletions were constructed with tet-sacB based selection and counterselection.^58^ C-terminal DAS+4 tag (with or without superfolder GFP tags) were added to chromosomal genes by direct integration and selected through integration of antibiotic resistance cassettes 3’ of the gene. All strains were confirmed by PCR, agarose gel electrophoresis and confirmed by sequencing (Eton Biosciences, or Genewiz) using paired oligonucleotides, either flanking the entire region. The recombineering plasmid pSIM5 and the tet-sacB selection/counterselection marker cassette were kind gifts from Donald Court (NCI, https://redrecombineering.ncifcrf.gov/court-lab.html). Strain BW25113 was obtained from the Yale Genetic Stock Center (CGSC http://cgsc.biology.yale.edu/). Strain DLF_R002 was constructed as previously reported by Menacho-Melgar et al.^32^ Strain DLFZ_0025 was constructed from DLF_R002 by first deleting the native *sspB* gene (using tet-sacB based selection and counterselection). Subsequently, the *cas3* gene was deleted^59^ and replaced with a low phosphate inducible sspB (using the *ugpB* gene promoter^28^) allele as well as a constitutive promoter to drive expression of the Cascade operon (again using tet-sacB based selection and counterselection). C-terminal DAS+4 tag modifications (with or without superfolder GFP tags) were added to the chromosome of DLF_Z0025 and its derivatives by direct integration and selected through integration of antibiotic resistance cassettes 3’ of the gene.

Plasmids, pCDF-ev (Addgene #89596), pHCKan-yibDp-GFPuv (Addgene #127078) and pHCKan-yibDp-cimA3.7 (Addgene #134595) were constructed as previously reported.^32^ Plasmids pCDF-mCherry1 (Addgene #87144) and pCDF-mCherry2 (Addgene #87145) were constructed from pCDF-ev by Gibson assembly of a PCR of the vector with synthetic DNA encoding an mCherry open reading frame with out without a C-terminal DAS+4 degron tag along with a strong synthetic constitutive proD promoter previously reported by Davis et al.^24^ pHCKan-ydbKp-sfGFP was constructed similarly.

Gene silencing guides and guide arrays were expressed from a series of pCASCADE plasmids. The pCASCADE-control plasmid was prepared by swapping the pTet promoter in pcrRNA.Tet (a kind gift from C. Beisel)^59^ with an insulated low phosphate induced ugpB promoter.^28^ In order to design CASCADE guide array, CASCADE PAM sites near the −35 or −10 box of the promoter of interest were identified, 30 bp at the 3’ end of PAM site was selected as the guide sequence and cloned into pCASCADE plasmid using Q5 site-directed mutagenesis (NEB, MA) following manufacturer’s protocol, with the modification that 5% v/v DMSO was added to the Q5 PCR reaction. PCR cycles were as follows: amplification involved an initial denaturation step at 98 °C for 30 second followed by cycling at 98 °C for 10 second, 72 °C for 30 second, and 72 °C for 1.5 min (the extension rate was 30 second/kb) for 25 cycles, then a final extension for 2 min at 72 °C. 2 μL of PCR mixture was used for 10 μL KLD reaction (NEB, MA), which proceeded under room temperature for 1 hour, after which, 1 μL KLD mixture was used for electroporation. The pCASCADE guide array plasmid (pCASCADE-G2Z) was prepared by sequentially amplifying complementary halves of each smaller guide plasmid by PCR, followed by subsequent DNA assembly as illustrated in Supplemental Materials. Primers used for pCASCADE assembly and gRNA sequences are provided in Supplemental Table S4. Additionally, all strains containing gRNA plasmids were routinely confirmed to assess gRNA stability via PCR as described below.

### BioLector studies

Single colonies of each strain were inoculated into 5 mL LB with appropriate antibiotics and cultured at 37 °C, 220 rpm for 9 hours or until OD600 reached > 2. 500 μL of the culture was inoculated into 10 mL SM10 medium with appropriate antibiotics, and cultured in a square shake flask (CAT#: 25-212, Genesee Scientific, Inc. San Diego, CA) at 37 °C, 220 rpm for 16 hours. Cells were pelleted by centrifugation and the culture density was normalized to OD600 = 5 using FGM3 media. Growth and fluorescence measurements were obtained in a Biolector (m2p labs,Baesweiler, Germany) using a high mass transfer FlowerPlate (CAT#: MTP-48-B, m2p-labs,Germany). 40 μL of the OD normalized culture was inoculated into 760 μL of FGM3 medium with appropriate antibiotics. Biolector settings were as follows: RFP gain=100, GFP gain=20, Biomass gain=20, shaking speed=1300 rpm, temperature=37 °C, humidity=85%. Every strain was analyzed in triplicate.

### ELISAs

Quantification of proteins via C-terminal GFP tags was performed using a GFP quantification kit from AbCam (Cambridge, UK, product # ab171581) according to manufacturer’s instructions. Briefly, samples were obtained from microfermentations as described above. Cells were harvested 24 hour post phosphate depletion, washed in water and lysed with the provided extraction buffer.

### NADPH Pools

NADPH was quantified with an NADPH Assay Kit (AbCam, Cambridge, UK, Cat # ab186031) according to manufacturer’s instructions. Briefly, cultures were grown in shake flasks (20mL fill in 250 mL Erlenmyer flasks) in SM10++ media and harvested in mid exponential phase, washed and resuspended in SM10 No phosphate media. After 24 hours of phosphate depletion, cells were pelleted by 10 minutes of centrifugation (4122 RCF, 4 °C) and lysed with the assay kit buffer. Cell debris was removed by centrifugation, generating cleared lysate used for quantification.

### Guide RNA Stability Testing

The stability of guide RNA arrays was confirmed by colony PCR using the following 2 primers: gRNA-for: 5’-GGGAGACCACAACGG-3’, gRNA-rev: 5’-CGCAGTCGAACGACCG-3’, using 2X EconoTaq Master mix (Lucigen) in 10 μL PCR reactions consisting of 5μL of 2X EconoTaq Master mix (Lucigen), 1uL of each primer (10μM), 3μL dH2O. A 98°C, 2-minute initial denaturation was followed by 35 cycles of 94°C, 30 seconds, 60°C 30 seconds, and 72°C, 30 seconds and a final 72°C, 5 min final extension. PCR reactions were then run on agarose gels and band size compared to control PCR reactions using purified plasmid DNA as a template. Guide protospacer loss occurred when guide array size was smaller than expected, indicating the loss of one or more protospacers.

### Fermentations

Minimal media microfermentations were performed as previously reported.^28^ 1L fermentations in instrumented bioreactors were also performed as previously reported,^60^ with slight modifications to the glucose feeding profiles, which were a function of strain and process. Generally, feeding was increased to enable excess residual glucose to ensure production rates were not feed limited. Glucose feeding was as follows. For 10 gCDW/L fermentations, starting batch glucose concentration was 25 g/L. A constant concentrated sterile filtered glucose feed (500 g/L) was added to the tanks at 1.5 g/h when cells entered mid-exponential growth. For 25 gCDW/L fermentations, starting batch glucose concentration was 25 g/L. Concentrated sterile filtered glucose feed (500 g/L) was added to the tanks at an initial rate of 7.1 g/h when cells entered mid-exponential growth. This rate was then increased exponentially, doubling every 1.083 hours (65 min) until 40 g total glucose had been added, after which the feed was maintained at 1.75g/hr.

### Production of Isotopically Labelled Metabolites

C^13^ pyruvate (CLM-1082-PK) and C^13^ D-glucose (U-13C6, 99%) were purchased from Cambridge Isotope Laboratories, Inc. (Tewksbury, MA). Isotopically labelled citramalate was produced in two stage minimal media shake flask studies, mimicking microfermentations, using strain DLF_Z0044 expressing cimA3.7. Briefly, 20 mL cultures of SM10++ media^60^ were inoculated with the strain which was grown overnight at 37 Celsius, shaking at 150 rpm in baffled 250 mL Erlenmyer shake flasks. After 16 hrs of growth cells were harvested by centrifugation washed and resuspended in 20mL of SM10 minimal media (lacking phosphate)^60^ where glucose was replaced with C^13^ labelled glucose. Cultures were grown for 25 hrs at 37 Celsius, shaking at 150 rpm, after which cells were removed by centrifugation, and the spent media filter sterilized prior to use as an internal standard.

### Analytical Methods

#### Cell dry weights

The OD/cell dry weight correlation coefficient (1 OD(600nm) = 0.35 gCDW/L, as determined by Menacho-Melgar et al. was used in this work.^60^

#### Glucose and Organic Acid Quantification

Two methods were used for glucose and organic acid quantification. First, a UPLC-RI method was developed for the simultaneous quantification of glucose, citramalate, acetic acid, pyruvate, citraconate, citrate and other organic acids including lactate, succinate, fumarate, malate, and mevalonate. Chromatographic separation was performed using a Rezex Fast Acid Analysis HPLC Column (100 x 7.8 mm, 9 μm particle size; CAT#: #1250100, Bio-Rad Laboratories, Inc., Hercules, CA) at 55 °C. 5 mM sulfuric acid was used as the isocratic eluent, with a flow rate of 0.5mL/min. Sample injection volume was 10 μL. Second, quantification was performed using a Bio-Rad Fast Acid Analysis HPLC Column (100 x 7.8 mm, 9 μm particle size; CAT#: #1250100, Bio-Rad Laboratories, Inc., Hercules, CA) at 65 °C. 10 mM sulfuric acid was used as the eluent, with an isocratic flow rate of 0.3mL/min. In both methods, sample injection volume was 10 μL and chromatography and detection were accomplished using a Waters Acquity H-Class UPLC integrated with a Waters 2414 Refractive Index (RI) detector (Waters Corp., Milford, MA. USA). Samples were diluted as needed to be within the accurate linear range. Dilution was performed using ultrapure water.

#### Organic acid Quantification via RapidFire-qTOF-MS

Micro-fermentation samples (as well as a confirmatory subset of samples from bioreactors) were centrifuged to remove cells. Broth was diluted 100 fold in water to a final volume of 20 μL. To this either a final concentration of 10 mg/L of C13 pyruvate was added or 2 uL of broth containing C13 labelled citramalate was added. The final sample was injected onto a HILIC (type H1 or the equivalent H6) RapidFire™ cartridge (Agilent Technologies, Santa Clara, CA). Injections were loaded on the cartridge with 95% hexane, 5% isopropanol for 3000 ms after a 600ms aspiration, at a flow rate of 1.0 mL/min. After loading, the cartridge was washed with isopropanol for 2000ms, at a flow rate of 1.0 mL/min. Elution was carried out for 8000 ms with 50% water/ 50% methanol with 0.2% acetic acid and 0.5 uM (NH_4_)_3_PO_4_,^61^ at a flow rate of 1.0 mL/min. Column equilibration was performed for 4000ms. The qTOF was tuned in the mass range of 50-250 m/z in fragile ion, negative ESI mode. Settings during detection were as follows: drying gas: 250C at a flow rate of 13L/minute, sheath gas: 400 C at a flow rate of 12L/minute, nebulizer pressure: 35 psi, Fragmenter voltage: 100 V, skimmer voltage: 65 V, nozzle voltage: 2000 V, capillary voltage: 3500V. The acquisition rate was 1 spectra/second.

## Acknowledgements

We would like to acknowledge the following support: NSF EAGER: #1445726, DARPA# HR0011-14-C-0075, ONR YIP #N00014-16-1-2558, DOE EERE grant #EE0007563 and financial support from DMC Biotechnologies, Inc. Additionally, we would also like to thank Michelle Luo and Chase Beisel for the kind gift of pcrRNA.Tet and support in implementation of Cascade based gene silencing.

## Author contributions

S. Li constructed plasmids and strains, performed microfermentations and fermentation studies and related analyses. Z. Ye constructed plasmids and strains, developed microfermetnation protocols, performed microfermentations and fermentation studies and related analyses. E.A. Moreb performed microfermentations and J. Lebeau performed ELISAs and fermentations. M.D. Lynch designed and analyzed experiments, constructed strains and performed ELISAs, and analytical analyses. All authors wrote revised and edited the manuscript.

## References

1. Bordbar, A., Monk, J. M., King, Z. A. & Palsson, B. O. Constraint-based models predict metabolic and associated cellular functions. Nat. Rev. Genet. 15, 107–120 (2014).

2. Burgard, A. P., Pharkya, P. & Maranas, C. D. Optknock: a bilevel programming framework for identifying gene knockout strategies for microbial strain optimization. Biotechnol. Bioeng. 84, 647–657 (2003).

3. Lynch, M. D. Into new territory: improved microbial synthesis through engineering of the essential metabolic network. Curr. Opin. Biotechnol. 38, 106–111 (2016).

4. Burg, J. M., Cooper, C. B., Ye, Z., Reed, B. R. & Moreb, E. A. Large-scale bioprocess competitiveness: the potential of dynamic metabolic control in two-stage fermentations. Current opinion in (2016).

5. Brockman, I. M. & Prather, K. L. J. Dynamic knockdown of E. coli central metabolism for redirecting fluxes of primary metabolites. Metab. Eng. 28, 104–113 (2015).

6. Brockman, I. M. & Prather, K. L. J. Dynamic metabolic engineering: New strategies for developing responsive cell factories. Biotechnol. J. 10, 1360–1369 (2015).

7. Venayak, N., von Kamp, A., Klamt, S. & Mahadevan, R. MoVE identifies metabolic valves to switch between phenotypic states. Nat. Commun. 9, 5332 (2018).

8. Solomon, K. V., Sanders, T. M. & Prather, K. L. J. A dynamic metabolite valve for the control of central carbon metabolism. Metabolic Engineering vol. 14 661–671 (2012).

9. Lalwani, M. A., Zhao, E. M. & Avalos, J. L. Current and future modalities of dynamic control in metabolic engineering. Curr. Opin. Biotechnol. 52, 56–65 (2018).

10. Gardner, T. S., Cantor, C. R. & Collins, J. J. Construction of a genetic toggle switch in Escherichia coli. Nature 403, 339–342 (2000).

11. David, F., Nielsen, J. & Siewers, V. Flux Control at the Malonyl-CoA Node through Hierarchical Dynamic Pathway Regulation in Saccharomyces cerevisiae. ACS Synth. Biol. 5, 224–233 (2016).

12. Xu, P., Li, L., Zhang, F., Stephanopoulos, G. & Koffas, M. Improving fatty acids production by engineering dynamic pathway regulation and metabolic control. Proc. Natl. Acad. Sci. U. S. A. 111, 11299–11304 (2014).

13. Kildegaard, K. R., Tramontin, L. R. R., Chekina, K. & Li, M. CRISPR/Cas9-RNA interference system for combinatorial metabolic engineering of Saccharomyces cerevisiae. Yeast (2019).

14. Ding, W., Zhang, Y. & Shi, S. Development and Application of CRISPR/Cas in Microbial Biotechnology. Frontiers in Bioengineering and Biotechnology 8, 711 (2020).

15. Qi, L. S. et al. Repurposing CRISPR as an RNA-guided platform for sequence-specific control of gene expression. Cell 152, 1173–1183 (2013).

16. Luo, M. L., Mullis, A. S., Leenay, R. T. & Beisel, C. L. Repurposing endogenous type I CRISPR-Cas systems for programmable gene repression. Nucleic Acids Research vol. 43 674–681 (2015).

17. McGinness, K. E., Baker, T. A. & Sauer, R. T. Engineering controllable protein degradation. Mol. Cell 22, 701–707 (2006).

18. Ye, Z., Lynch, M.D., Trahan, A.D., Rodriguez, D.L., Cooper, C.B. Bozdag, A. Compositions and methods for rapid and dynamic flux control using synthetic metabolic valves. Patent (2015).

19. Menacho-Melgar, R., Moreb, E. A. & Efromson, J. P. Improved, two-stage protein expression and purification via autoinduction of both autolysis and auto DNA/RNA hydrolysis conferred by phage lysozyme and DNA/RNA…. bioRxiv (2020).

20. Huber, R., Roth, S., Rahmen, N. & Buchs, J. Utilizing high-throughput experimentation to enhance specific productivity of an E.coli T7 expression system by phosphate limitation. BMC Biotechnol. 11, 22 (2011).

21. Chubukov, V. & Sauer, U. Environmental dependence of stationary-phase metabolism in Bacillus subtilis and Escherichia coli. Appl. Environ. Microbiol. 80, 2901–2909 (2014).

22. Menacho-Melgar, R. et al. Improved two-stage protein expression and purification via autoinduction of both autolysis and auto DNA/RNA hydrolysis conferred by phage lysozyme and DNA/RNA endonuclease. Biotechnol. Bioeng. (2020) doi:10.1002/bit.27444.

23. Brouns, S. J. J. et al. Small CRISPR RNAs guide antiviral defense in prokaryotes. Science 321, 960–964 (2008).

24. Davis, J. H., Rubin, A. J. & Sauer, R. T. Design, construction and characterization of a set of insulated bacterial promoters. Nucleic Acids Res. 39, 1131–1141 (2011).

25. Hintsche, M. & Klumpp, S. Dilution and the theoretical description of growth-rate dependent gene expression. J. Biol. Eng. 7, 22 (2013).

26. Grünenfelder, B. et al. Proteomic analysis of the bacterial cell cycle. Proc. Natl. Acad. Sci. U. S. A. 98, 4681–4686 (2001).

27. Pédelacq, J.-D., Cabantous, S., Tran, T., Terwilliger, T. C. & Waldo, G. S. Engineering and characterization of a superfolder green fluorescent protein. Nat. Biotechnol. 24, 79–88 (2006).

28. Moreb, E. A. et al. Media Robustness and scalability of phosphate regulated promoters useful for two-stage autoinduction in E. coli. ACS Synthetic Biology (2020) doi:10.1021/acssynbio.0c00182.

29. Huergo, L. F. & Dixon, R. The Emergence of 2-Oxoglutarate as a Master Regulator Metabolite. Microbiol. Mol. Biol. Rev. 79, 419–435 (2015).

30. Rabinowitz, J. D. & Silhavy, T. J. Systems biology: metabolite turns master regulator. Nature vol. 500 283–284 (2013).

31. Doucette, C. D., Schwab, D. J., Wingreen, N. S. & Rabinowitz, J. D. α-Ketoglutarate coordinates carbon and nitrogen utilization via enzyme I inhibition. Nat. Chem. Biol. 7, 894–901 (2011).

32. Menacho-Melgar, R. et al. Improved, scalable, two-stage, autoinduction of recombinant protein expression in E. coli utilizing phosphate depletion. doi:10.1101/820787.

33. Moreb, E. A. et al. Robustness testing and scalability of phosphate regulated promoters useful for two-stage autoinduction in E. coli. (2020) doi:10.1021/acssynbio.0c00182.

34. Lebeau, J., Efromson, J. P. & Lynch, M. D. A Review of the Biotechnological Production of Methacrylic Acid. Frontiers in Bioengineering and Biotechnology 8, 207 (2020).

35. Howell, D. M., Xu, H. & White, R. H. (R)-citramalate synthase in methanogenic archaea. J. Bacteriol. 181, 331–333 (1999).

36. Parimi, N. S., Durie, I. A., Wu, X., Niyas, A. M. M. & Eiteman, M. A. Eliminating acetate formation improves citramalate production by metabolically engineered Escherichia coli. Microb. Cell Fact. 16, 114 (2017).

37. Wu, X. & Eiteman, M. A. Production of citramalate by metabolically engineered Escherichia coli. Biotechnol. Bioeng. 113, 2670–2675 (2016).

38. Webb, J. P. et al. Efficient bio-production of citramalate using an engineered Escherichia coli strain. Microbiology 164, 133–141 (2018).

39. Webb, J. et al. Systems Analyses Reveal the Resilience of Escherichia coli Physiology during Accumulation and Export of the Nonnative Organic Acid Citramalate. mSystems 4, (2019).

40. Atsumi, S. & Liao, J. C. Directed evolution of Methanococcus jannaschii citramalate synthase for biosynthesis of 1-propanol and 1-butanol by Escherichia coli. Appl. Environ. Microbiol. 74, 7802–7808 (2008).

41. de Kok, A., Hengeveld, A. F., Martin, A. & Westphal, A. H. The pyruvate dehydrogenase multi-enzyme complex from Gram-negative bacteria. Biochim. Biophys. Acta 1385, 353–366 (1998).

42. Hennig, J. et al. Molecular mechanism of regulation of the pyruvate dehydrogenase complex from E. coli. Biochemistry 36, 15772–15779 (1997).

43. Griffith, K. L. & Wolf, R. E. Systematic mutagenesis of the DNA binding sites for SoxS in the Escherichia coli zwf and fpr promoters: identifying nucleotides required for DNA binding and transcription activation. Molecular Microbiology vol. 42 571–571 (2001).

44. Greenberg, J. T., Monach, P., Chou, J. H., Josephy, P. D. & Demple, B. Positive control of a global antioxidant defense regulon activated by superoxide-generating agents in Escherichia coli. Proc. Natl. Acad. Sci. U. S. A. 87, 6181–6185 (1990).

45. Christodoulou, D. et al. Reserve Flux Capacity in the Pentose Phosphate Pathway Enables Escherichia coli’s Rapid Response to Oxidative Stress. Cell Syst 6, 569–578.e7 (2018).

46. Nakayama, T., Yonekura, S.-I., Yonei, S. & Zhang-Akiyama, Q.-M. Escherichia coli pyruvate:flavodoxin oxidoreductase, YdbK - regulation of expression and biological roles in protection against oxidative stress. Genes & Genetic Systems vol. 88 175–188 (2013).

47. Blaschkowski, H. P., Knappe, J., Ludwig-Festl, M. & Neuer, G. Routes of Flavodoxin and Ferredoxin Reduction in Escherichia coli: CoA-Acylating Pyruvate: Flavodoxin and NADPH: Flavodoxin Oxidoreductases Participating in the Activation of Pyruvate Formate-Lyase. Eur. J. Biochem. 123, 563–569 (1982).

48. Akhtar, M. K., Kalim Akhtar, M. & Jones, P. R. Construction of a synthetic YdbK-dependent pyruvate:H2 pathway in Escherichia coli BL21(DE3). Metabolic Engineering vol. 11 139–147 (2009).

49. Fàbrega, A., Rosner, J. L., Martin, R. G., Solé, M. & Vila, J. SoxS-dependent coregulation of ompN and ydbK in a multidrug-resistant Escherichia coli strain. FEMS Microbiol. Lett. 332, 61–67 (2012).

50. Koo, M.-S. et al. A reducing system of the superoxide sensor SoxR in Escherichia coli. EMBO J. 22, 2614–2622 (2003).

51. Gaudu, P., Dubrac, S. & Touati, D. Activation of SoxR by Overproduction of Desulfoferrodoxin: Multiple Ways To Induce the soxRSRegulon. J. Bacteriol. 182, 1761–1763 (2000).

52. Giró, M., Carrillo, N. & Krapp, A. R. Glucose-6-phosphate dehydrogenase and ferredoxin-NADP(H) reductase contribute to damage repair during the soxRS response of Escherichia coli. Microbiology vol. 152 1119–1128 (2006).

53. Jair, K.-W., Fawcett, W. P., Fujita, N., Ishihama, A. & Wolf, R. E., Jr. Ambidextrous transcriptional activation by SoxS: requirement for the C-terminal domain of the RNA polymerase alpha subunit in a subset of Escherichia coli superoxide-inducible genes. Mol. Microbiol. 19, 307–317 (1996).

54. Bianchi, V., Haggård-Ljungquist, E., Pontis, E. & Reichard, P. Interruption of the ferredoxin (flavodoxin) NADP+ oxidoreductase gene of Escherichia coli does not affect anaerobic growth but increases sensitivity to paraquat. J.Bacteriol. 177, 4528–4531 (1995).

55. Maury, J. et al. Glucose-Dependent Promoters for Dynamic Regulation of Metabolic Pathways. Front Bioeng Biotechnol 6, 63 (2018).

56. Soma, Y., Tsuruno, K., Wada, M., Yokota, A. & Hanai, T. Metabolic flux redirection from a central metabolic pathway toward a synthetic pathway using a metabolic toggle switch. Metab. Eng. 23, 175–184 (2014).

57. Eremina, N. S. Overexpression of ydbKencoding putative pyruvate synthase improves L-valine production and aerobic growth on ethanol media by an Escherichia coli strain carrying an oxygen-resistant alcohol dehydrogenase. J. Microbial Biochem. Technol. 2, 77–83 (2010).

58. Li, X.-T., Thomason, L. C., Sawitzke, J. A., Costantino, N. & Court, D. L. Positive and negative selection using the tetA-sacB cassette: recombineering and P1 transduction in Escherichia coli. Nucleic Acids Res. 41, e204 (2013).

59. Luo, M. L., Mullis, A. S., Leenay, R. T. & Beisel, C. L. Repurposing endogenous type I CRISPR-Cas systems for programmable gene repression. NucleicAcidsRes. 43, 674–681 (2015).

60. Menacho-Melgar, R. et al. Scalable, two-stage, autoinduction of recombinant protein expression in E. coli utilizing phosphate depletion. Biotechnol. Bioeng. 26, 44 (2020).

61. Spalding, J. L., Naser, F. J., Mahieu, N. G., Johnson, S. L. & Patti, G. J. Trace Phosphate Improves ZIC-pHILIC Peak Shape, Sensitivity, and Coverage for Untargeted Metabolomics. J. Proteome Res. 17, 3537–3546 (2018).

