## Supplementary Materials for "Dynamic control over feedback regulation identifies pyruvate-ferredoxin oxidoreductase as a central metabolic enzyme in stationary phase *E. coli*"

### Supplemental Materials

#### Section 1: Modified Strains

**Table S1:** List of chromosomally modified strains.

| Strain | Genotype | Source |
|---|---|---|
| DLF_R002 | F-, λ-, Δ(araD-araB)567, lacZ4787(del)(::rrnB-3) , rph-1, Δ(rhaD-rhaB)568, hsdR514, ΔackA-pta, ΔpoxB, ΔpflB, ΔldhA, ΔadhE, ΔiclR, ΔarcA | [1] |
| DLF_R002 | DLF_R002, ΔsspB | this study |
| DLF_Z002 | DLF_Z002, Δcas3::tm-ugpb-sspB-pro-casA | this study |
| DLF_Z01517 | DLF_Z002, Δcas3::tm-pro-casA | this study |
| DLF_Z0043 | DLF_Z0025, gltA-DAS+4-zeoR | this study |
| DLF_Z0043G | DLF_Z0025, gltA-sfGFP-zeoR | this study |
| DLF_Z0043GD | DLF_Z0025, gltA-sfGFP-DAS+4-zeoR | this study |
| DLF_Z01002 | DLF_Z0025, zwf-DAS+4-bsdR | this study |
| DLF_Z01002G | DLF_Z0025, zwf-sfGFP-zeoR | this study |
| DLF_Z01002GD | DLF_Z0025, zwf-sfGFP-DAS+4-zeoR | this study |
| DLF_Z0044 | DLF_Z0025, gltA-DAS+4-zeoR, zwf-DAS+4-bsdR | this study |
| DLF_Z0048 | DLF_Z0025, lpd-DAS+4-gentR, gltA-DAS+4-zeoR, zwf-DAS+4-bsdR | this study |

**Table S2:** Oligonucleotides utilized for strain construction.

| Oligo | Sequence |
|---|---|
| SL1 | CAG TCC AGT TAC GCT GGA GTC |
| SR2 | GGT CAG GTA TGA TTT AA A TGG TCA GT |
| sspB_kan_F | CTGGTACACGCTGATGAACACC |
| sspB_kan_R | CTGGTCATTGCCATTTGTGCC |
| sspB_conf_F | GAATCAGAGCGTTCCGACCC |
| sspB_conf_R | GTACGCAGTTTGCCAACGTG |
| cas3_tetA_F | AATAGCCCGCTGATATCATCGATAATACTAAAAAAACAGGGAGGCTATT ATCCTAATTTTTGTTGACACTCTATC |

| | |
|---|---|
| cas3_sacB_R | TACAGGGATCCAGTTATCAATAAGCAAATTCATTTGTTCTCCTTCATATG ATCAAAGGGAAAACTGTCCATATGC |
| cas3_conf_F | CAAGACATGTGTATATCACTGTAATTC |
| cas3_500dn | GCGATTGCAGATTTATGATTTGG |
| gltA_conf_F | TATCATCCTGAAAGCGATGG |
| zwf_conf_F | CTGCTGGAAACCATGCG |
| bsdR_intR | GAGCATGGTGATCTTCTCAGT |
| zeoR_intR | ACTGAAGCCCAGACGATC |
| lpd_conf_F | ATCTCACCGTGTGATCGG |
| gentR_intR | GCGATGAATGTCTTACTACGGA |
| yibDpF | GTGCGTAATTGTGCTGATC |
| yibDpF_rc | GATCAGCACAATTACGCAC |
| pCOLA-Amp1 | ACTGACCATTTAAATCATACCTGACCTAGagagtcacactggctc |

**Table S3:** Synthetic DNA utilized for strain construction.

| **tetA-sacB Cassette** |
|---|
| TCCTAATTTTTGTTGACACTCTATCATTGATAGAGTTATTTTACCACTCCCTATCAGTGATA GAGAAAAGTGAAATGAATAGTTCGACAAAGATCGCATTGGTAATTACGTTACTCGATGCC ATGGGGATTGGCCTTATCATGCCAGTCTTGCCAACGTTATTACGTGAATTTATTGCTTCGG AAGATATCGCTAACCACTTTGGCGTATTGCTTGCACTTTATGCGTTAATGCAGGTTATCTTT GCTCCTTGGCTTGGAAAAATGTCTGACCGATTTGGTCGGCGCCCAGTGCTGTTGTTGTCAT TAATAGGCGCATCGCTGGATTACTTATTGCTGGCTTTTTCAAGTGCGCTTTGGATGCTGTAT TTAGGCCGTTTGCTTTCAGGGATCACAGGAGCTACTGGGGCTGTCGCGGCATCGGTCATTG CCGATACCACCTCAGCTTCTCAACGCGTGAAGTGGTTCGGTTGGTTAGGGGCAAGTTTTGG GCTTGGTTTAATAGCGGGGCCTATTATTGGTGGTTTTGCAGGAGAGATTTCACCGCATAGT CCCTTTTTTATCGCTGCGTTGCTAAATATTGTCACTTTCCTTGTGGTTATGTTTTGGTTCCGT GAAACCAAAAATACACGTGATAATACAGATACCGAAGTAGGGGTTGAGACGCAATCGAA TTCGGTATACATCACTTTATTTAAAACGATGCCCATTTTGTTGATTATTTATTTTTCAGCGC AATTGATAGGCCAAATTCCCGCAACGGTGTGGGTGCTATTTACCGAAAATCGTTTTGGATG GAATAGCATGATGGTTGGCTTTTCATTAGCGGGTCTTGGTCTTTTACACTCAGTATTCCAA GCCTTTGTGGCAGGAAGAATAGCCACTAAATGGGGCGAAAAAACGGCAGTACTGCTCGG ATTTATTGCAGATAGTAGTGCATTTGCCTTTTTAGCGTTTATATCTGAAGGTTGGTTAGTTT TCCCTGTTTTAATTTTATTGGCTGGTGGTGGGATCGCTTTACCTGCATTACAGGGAGTGATG TCTATCCAAACAAAGAGTCATCAGCAAGGTGCTTTACAGGGATTATTGGTGAGCCTTACC AATGCAACCGGTGTTATTGGCCCATTACTGTTTGCTGTTATTTATAATCATTCACTACCAAT TTGGGATGGCTGGATTTGGATTATTGGTTTAGCGTTTTACTGTATTATTATCCTGCTATCGA TGACCTTCATGTTAACCCCTCAAGCTCAGGGGAGTAAACAGGAGACAAGTGCTTAGTTAT TTCGTCACCAAATGATGTTATTCCGCGAAATATAATGACCCTCTTGATAACCCAAGAGCAT CACATATACCTGCCGTTCACTATTATTTAGTGAAATGAGATATTATGATATTTCTGAATTG TGATTAAAAAGGCAACTTTATGCCCATGCAACAGAAACTATAAAAAATACAGAGAATGAA |

| |
|---|
| AAGAAACAGATAGATTTTTTAGTTCTTTAGGCCCGTAGTCTGCAAATCCTTTTATGATTTTC<br>TATCAAACAAAAGAGGAAAATAGACCAGTTGCAATCCAAACGAGAGTCTAATAGAATGA<br>GGTCGAAAAGTAAATCGCGCGGGTTTGTTACTGATAAAGCAGGCAAGACCTAAAATGTGT<br>AAAGGGCAAAGTGTATACTTTGGCGTCACCCCTTACATATTTTAGGTCTTTTTTTATTGTGC<br>GTAACTAACTTGCCATCTTCAAACAGGAGGGCTGGAAGAAGCAGACCGCTAACACAGTAC<br>ATAAAAAAGGAGACATGAACGATGAACATCAAAAAGTTTGCAAAACAAGCAACAGTATT<br>AACCTTTACTACCGCACTGCTGGCAGGAGGCGCAACTCAAGCGTTTGCGAAAGAAACGAA<br>CCAAAAGCCATATAAGGAAACATACGGCATTTCCCATATTACACGCCATGATATGCTGCA<br>AATCCCTGAACAGCAAAAAAAATGAAAAATATCAAGTTCCTGAGTTCGATTCGTCCACAAT<br>TAAAAATATCTCTTCTGCAAAAGGCCTGGACGTTTGGGACAGCTGGCCATTACAAAACGC<br>TGACGGCACTGTCGCAAACTATCACGGCTACCACATCGTCTTTGCATTAGCCGGAGATCCT<br>AAAAATGCGGATGACACATCGATTTACATGTTCTATCAAAAAGTCGGCGAAACTTCTATT<br>GACAGCTGGAAAAACGCTGGCCGCGTCTTTAAAGACAGCGACAAATTCGATGCAAATGAT<br>TCTATCCTAAAAGACCAAACACAAGAATGGTCAGGTTCAGCCACATTTACATCTGACGGA<br>AAAATCCGTTTATTCTACACTGATTTCTCCGGTAAACATTACGGCAAACAAACACTGACAA<br>CTGCACAAGTTAACGTATCAGCATCAGACAGCTCTTTGAACATCAACGGTGTAGAGGATT<br>ATAAATCAATCTTTGACGGTGACGGAAAAACGTATCAAAATGTACAGCAGTTCATCGATG<br>AAGGCAACTACAGCTCAGGCGACAACCATACGCTGAGAGATCCTCACTACGTAGAAGATA<br>AAGGCCACAAATACTTAGTATTTGAAGCAAACACTGGAACTGAAGATGGCTACCAAGGCG<br>AAGAATCTTTATTTAACAAAGCATACTATGGCAAAAGCACATCATTCTTCCGTCAAGAAA<br>GTCAAAAACTTCTGCAAAGCGATAAAAAACGCACGGCTGAGTTAGCAAACGGCGCTCTCG<br>GTATGATTGAGCTAAACGATGATTACACACTGAAAAAAGTGATGAAACCGCTGATTGCAT<br>CTAACACAGTAACAGATGAAATTGAACGCGCGAACGTCTTTAAAATGAACGGCAAATGGT<br>ACCTGTTCACTGACTCCCGCGGATCAAAAATGACGATTGACGGCATTACGTCTAACGATAT<br>TTACATGCTTGGTTATGTTTCTAATTCTTTAACTGGCCCATACAAGCCGCTGAACAAAACT<br>GGCCTTGTGTTAAAAATGGATCTTGATCCTAACGATGTAACCTTTACTTACTCACACTTCG<br>CTGTACCTCAAGCGAAAGGAAACAATGTCGTGATTACAAGCTATATGACAAACAGAGGAT<br>TCTACGCAGACAAACAATCAACGTTTGCGCCAAGCTTCCTGCTGAACATCAAAGGCAAGA<br>AAACATCTGTTGTCAAAGACAGCATCCTTGAACAAGGACAATTAACAGTTAACAAATAAA<br>AACGCAAAAGAAAATGCCGATATTGACTACCGGAAGCAGTGTGACCGTGTGCTTCTCAAA<br>TGCCTGATTCAGGCTGTCTATGTGTGACTGTTGAGCTGTAACAAGTTGTCTCAGGTGTTCA<br>ATTTCATGTTCTAGTTGCTTTGTTTTACTGGTTTCACCTGTTCTATTAGGTGTTACATGCTGT<br>TCATCTGTTACATTGTCGATCTGTTCATGGTGAACAGCTTTAAATGCACCAAAAACTCGTA<br>AAAGCTCTGATGTATCTATCTTTTTTACACCGTTTTCATCTGTGCATATGGACAGTTTTCCC<br>TTTGAT |
| **Δcas3-pro-casA** |
| CAAGACATGTGTATATCACTGTAATTCGATATTTATGAGCAGCATCGAAAAATAGCCCGCT<br>GATATCATCGATAATACTAAAAAAACAGGGAGGCTATTACCAGGCATCAAATAAAACGA<br>AAGGCTCAGTCGAAAGACTGGGCCTTTCGTTTTATCTGTTGTTTGTCGGTGAACGCTCTCT<br>ACTAGAGTCACACTGGCTCACCTTCGGGTGGGCCTTTCTGCGTTTATATCTTTCTGACACCT<br>TACTATCTTACAAATGTAACAAAAAAGTTATTTTTCTGTAATTCGAGCATGTCATGTTACC<br>CCGCGAGCATAAAACGCGTGTGTAGGAGGATAATCTTTGACGGCTAGCTCAGTCCTAGGT<br>ACAGTGCTAGCCATATGAAGGAGAACAAATGAATTTGCTTATTGATAACTGGATCCCTGT<br>ACGCCCGCGAAACGGGGGGAAAGTCCAAATCATAAATCTGCAATCGCTATAC |
| **Δcas3::ugBp-sspB-pro-casA** |
| CAAGACATGTGTATATCACTGTAATTCGATATTTATGAGCAGCATCGAAAAATAGCCCGCT<br>GATATCATCGATAATACTAAAAAAACAGGGAGGCTATTACCAGGCATCAAATAAAACGA |

| |
|---|
| AAGGCTCAGTCGAAAGACTGGGCCTTTCGTTTTATCTGTTGTTTGTCGGTGAACGCTCTCT<br>ACTAGAGTCACACTGGCTCACCTTCGGGTGGGCCTTTCTGCGTTTATATCTTTCTGACACCT<br>TACTATCTTACAAATGTAACAAAAAAGTTATTTTTCTGTAATTCGAGCATGTCATGTTACC<br>CCGCGAGCATAAAACGCGTGTGTAGGAGGATAATCTATGGATTTGTCACAGCTAACACCA<br>CGTCGTCCCTATCTGCTGCGTGCATTCTATGAGTGGTTGCTGGATAACCAGCTCACGCCGC<br>ACCTGGTGGTGGATGTGACGCTCCCTGGCGTGCAGGTTCCTATGGAATATGCGCGTGACG<br>GGCAAATCGTACTCAACATTGCGCCGCGTGCTGTCGGCAATCTGGAACTGGCGAATGATG<br>AGGTGCGCTTTAACGCGCGCTTTGGTGGCATTCCGCGTCAGGTTTCTGTGCCGCTGGCTGC<br>CGTGCTGGCTATCTACGCCCGTGAAAATGGCGCAGGCACGATGTTTGAGCCTGAAGCTGC<br>CTACGATGAAGATACCAGCATCATGAATGATGAAGAGGCATCGGCAGACAACGAAACCG<br>TTATGTCGGTTATTGATGGCGACAAGCCAGATCACGATGATGACACTCATCCTGACGATG<br>AACCTCCGCAGCCACCACGCGGTGGTCGACCGGCATTACGCGTTGTGAAGTAATTGACGG<br>CTAGCTCAGTCCTAGGTACAGTGCTAGCCATATGAAGGAGAACAAATGAATTTGCTTATT<br>GATAACTGGATCCCTGTACGCCCGCGAAACGGGGGGAAAGTCCAAATCATAAATCTGCAA<br>TCGCTATAC |
| **gltA-DAS+4-zeoR** |
| GTATTCCGTCTTCCATGTTCACCGTCATTTTCGCAATGGCACGTACCGTTGGCTGGATCGCC<br>CACTGGAGCGAAATGCACAGTGACGGTATGAAGATTGCCCGTCCGCGTCAGCTGTATACA<br>GGATATGAAAAACGCGACTTTAAAAGCGATATCAAGCGTGCGGCCAACGATGAAAACTAT<br>TCTGAAAACTATGCGGATGCGTCTTAATAGTTGACAATTAATCATCGGCATAGTATATCGG<br>CATAGTATAATACGACTCACTATAGGAGGGCCATCATGGCCAAGTTGACCAGTGCCGTTC<br>CGGTGCTCACCGCGCGCGACGTCGCCGGAGCGGTCGAGTTCTGGACCGACCGGCTCGGGT<br>TCTCCCGGGACTTCGTGGAGGACGACTTCGCCGGTGTGGTCCGGGACGACGTGACCCTGTT<br>CATCAGCGCGGTCCAGGACCAGGTGGTGCCGGACAACACCCTGGCCTGGGTGTGGGTGCG<br>CGGCCTGGACGAGCTGTACGCCGAGTGGTCGGAGGTCGTGTCCACGAACTTCCGGGACGC<br>CTCCGGGCCGGCCATGACCGAGATCGGCGAGCAGCCGTGGGGGCGGGAGTTCGCCCTGCG<br>CGACCCGGCCGGCAACTGCGTGCACTTTGTGGCAGAGGAGCAGGACTGAGGATAAGTAAT<br>GGTTGATTGCTAAGTTGTAAATATTTTAACCCGCCGTTCATATGGCGGGTTGATTTTTATAT<br>GCCTAAACACAAAAAATTGTAAAAATAAAATCCATTAACAGACCTATATAGATATTTAAA<br>AAGAATAGAACAGCTCAAATTATCAGCAACCCAATACTTTCAATTAAAAACTTCATGGTA<br>GTCGCATTTATAACCCTATGAAA |
| **gltA-sfGFP-zeoR** |
| AACGTCGATTTCTACTCTGGTATCATCCTGAAAGCGATGGGTATTCCGTCTTCCATGTTCA<br>CCGTCATTTTCGCAATGGCACGTACCGTTGGCTGGATCGCCCACTGGAGCGAAATGCACA<br>GTGACGGTATGAAGATTGCCCGTCCGCGTCAGCTGTATACAGGATATGAAAAACGCGACT<br>TTAAAAGCGATATCAAGCGTGGGGGTTCAGGCGGGTCGGGTGGCgtgagcaagggcgaggagctgttca<br>ccggggtggtgcccatcctggtcgagctggacggcgacgtaaacggccacaagttcagcgtgcgcggcgagggcgagggcgatgccaccaacg<br>gcaagctgaccctgaagttcatctgcaccaccggcaagctgcccgtgccctggcccaccctcgtgaccaccctgacctacggcgtgcagtgcttca<br>gccgctaccccgaccacatgaagcgccacgacttcttcaagtccgccatgcccgaaggctacgtccaggagcgcaccatcagcttcaaggacgac<br>ggcacctacaagacccgcgccgaggtgaagttcgagggcgacaccctggtgaaccgcatcgagctgaagggcatcgacttcaaggaggacggc<br>aacatcctggggcacaagctggagtacaacttcaacagccacaacgtctatatcaccgccgacaagcagaagaacggcatcaaggccaacttcaa<br>gatccgccacaacgtggaggacggcagcgtgcagctcgccgaccactaccagcagaacacccccatcggcgacggccccgtgctgctgcccga<br>caaccactacctgagcacccagtccgtgctgagcaaagaccccaacgagaagcgcgatcacatggtcctgctggagttcgtgaccgccgccggga<br>tcactcacggcatggacgagctgtacaagTAATGATGATCGGCACGTAAGAGGTTCCAACTTTCACCATAA<br>TGAAATAAGATCACTACCGGGCGTATTTTTTGAGTTATCGAGATTTTCAGGAGCTAAGGAA<br>GCTAAAATGGCTAAACTGACGTCGGCCGTTCCAGTGCTTACTGCGCGTGATGTAGCGGGA<br>GCCGTAGAGTTTTGGACGGATCGTCTTGGGTTTAGTCGCGACTTTGTGGAAGATGACTTCG |

| |
|---|
| CAGGGGTTGTTCGTGATGACGTCACACTGTTCATCAGTGCCGTACAGGATCAGGTTGTACC<br>CGATAACACTCTTGCGTGGGTATGGGTGCGTGGCCTGGATGAGTTATACGCCGAATGGTC<br>CGAGGTAGTCAGCACAAACTTCCGCGACGCATCCGGGCCCGCTATGACTGAGATCGGGGA<br>ACAACCGTGGGGACGTGAGTTTGCCTTACGTGACCCGGCGGGGAACTGCGTCCACTTTGT<br>GGCGGAGGAGCAGGACTAAGGATAAGtagTGGTTGATTGCTAAGTTGTAAATATTTTAACC<br>CGCCGTTCATATGGCGGGTTGATTTTTATATGCCTAAACACAAAAAATTGTAAAAATAAAA<br>TCCATTAACAGACCTATATAGATATTTAAAAAGAATAGAACAGCTCAAATTATCAGCAAC<br>CCAATACTTTCAATTAAAAACTTCATGGTAGTCGCATTTATAACCCTATGAAAATGACGTC<br>TATCTATACCCCCCTATATTTTATTCATCATACAACAAATTCATGATACCAATAA |
| **gltA-sfGFP-DAS+4-zeoR** |
| AACGTCGATTTCTACTCTGGTATCATCCTGAAAGCGATGGGTATTCCGTCTTCCATGTTCA<br>CCGTCATTTTCGCAATGGCACGTACCGTTGGCTGGATCGCCCACTGGAGCGAAATGCACA<br>GTGACGGTATGAAGATTGCCCGTCCGCGTCAGCTGTATACAGGATATGAAAAACGCGACT<br>TTAAAAGCGATATCAAGCGTGGGGGTTCAGGCGGGTCGGGTGGCgtgagcaagggcgaggagctgttca<br>ccggggtggtgcccatcctggtcgagctggacggcgacgtaaacggccacaagttcagcgtgcgcggcgagggcgagggcgatgccaccaacg<br>gcaagctgaccctgaagttcatctgcaccaccggcaagctgcccgtgccctggcccaccctcgtgaccaccctgacctacggcgtgcagtgcttca<br>gccgctaccccgaccacatgaagcgccacgacttcttcaagtccgccatgcccgaaggctacgtccaggagcgcaccatcagcttcaaggacgac<br>ggcacctacaagacccgcgccgaggtgaagttcgagggcgacaccctggtgaaccgcatcgagctgaagggcatcgacttcaaggaggacggc<br>aacatcctggggcacaagctggagtacaacttcaacagccacaacgtctatatcaccgccgacaagcagaagaacggcatcaaggccaacttcaa<br>gatccgccacaacgtggaggacggcagcgtgcagctcgccgaccactaccagcagaacacccccatcggcgacggccccgtgctgctgcccga<br>caaccactacctgagcacccagtccgtgctgagcaaagaccccaacgagaagcgcgatcacatggtcctgctggagttcgtgaccgccgccggga<br>tcactcacggcatggacgagctgtacaagGGTGGGGGTGGAGCGGCGGCGGTGGCTCCGCGGCCAACGA<br>TGAAAACTATTCTGAAAACTATGCGGATGCGTCTTAATGATGATCGGCACGTAAGAGGTT<br>CCAACTTTCACCATAATGAAATAAGATCACTACCGGGCGTATTTTTTGAGTTATCGAGATT<br>TTCAGGAGCTAAGGAAGCTAAAATGGCTAAACTGACGTCGGCCGTTCCAGTGCTTACTGC<br>GCGTGATGTAGCGGGAGCCGTAGAGTTTTGGACGGATCGTCTTGGGTTTAGTCGCGACTTT<br>GTGGAAGATGACTTCGCAGGGGTTGTTCGTGATGACGTCACACTGTTCATCAGTGCCGTAC<br>AGGATCAGGTTGTACCCGATAACACTCTTGCGTGGGTATGGGTGCGTGGCCTGGATGAGT<br>TATACGCCGAATGGTCCGAGGTAGTCAGCACAAACTTCCGCGACGCATCCGGGCCCGCTA<br>TGACTGAGATCGGGGAACAACCGTGGGGACGTGAGTTTGCCTTACGTGACCCGGCGGGGA<br>ACTGCGTCCACTTTGTGGCGGAGGAGCAGGACTAAGGATAAGtagTGGTTGATTGCTAAGTT<br>GTAAATATTTTAACCCGCCGTTCATATGGCGGGTTGATTTTTATATGCCTAAACACAAAAA<br>ATTGTAAAAATAAAATCCATTAACAGACCTATATAGATATTTAAAAAGAATAGAACAGCT<br>CAAATTATCAGCAACCCAATACTTTCAATTAAAAACTTCATGGTAGTCGCATTTATAACCC<br>TATGAAAATGACGTCTATCTATACCCCCCTATATTTTATTCATCATACAACAAATTCATGA<br>TACCAATAA |
| **zwf-DAS+4-bsdR** |
| GAAGTGGAAGAAGCCTGGAAATGGGTAGACTCCATTACTGAGGCGTGGGCGATGGACAA<br>TGATGCGCCGAAACCGTATCAGGCCGGAACCTGGGGACCCGTTGCCTCGGTGGCGATGAT<br>TACCCGTGATGGTCGTTCCTGGAATGAGTTTGAGGCGGCCAACGATGAAAACTATTCTGA<br>AAACTATGCGGATGCGTCTTAATAGTTGACAATTAATCATCGGCATAGTATATCGGCATAG<br>TATAATACGACTCACTATAGGAGGGCCATCATGAAGACCTTCAACATCTCTCAGCAGGAT<br>CTGGAGCTGGTGGAGGTCGCCACTGAGAAGATCACCATGCTCTATGAGGACAACAAGCAC<br>CATGTCGGGGCGGCCATCAGGACCAAGACTGGGGAGATCATCTCTGCTGTCCACATTGAG<br>GCCTACATTGGCAGGGTCACTGTCTGTGCTGAAGCCATTGCCATTGGGTCTGCTGTGAGCA<br>ACGGGCAGAAGGACTTTGACACCATTGTGGCTGTCAGGCACCCCTACTCTGATGAGGTGG<br>ACAGATCCATCAGGGTGGTCAGCCCCTGTGGCATGTGCAGAGAGCTCATCTCTGACTATG |

| |
|---|
| CTCCTGACTGCTTTGTGCTCATTGAGATGAATGGCAAGCTGGTCAAAACCACCATTGAGGA ACTCATCCCCCTCAAGTACACCAGGAACTAAAGTAATATCTGCGCTTATCCTTTATGGTTA TTTTACCGGTAACATGATCTTGCGCAGATTGTAGAACAATTTTTACACTTTCAGGCCTCGT GCGGATTCACCCAC GAGGCTTTTTTTATTACACTGACTGAAACGTTTTTGCCCTATGAGCTCCGGTTACAGGCGTT TCAGTCATAAATCCTCTGAATGAAACGCGTTGTGAATC |
| **zwf-sfGFP-zeoR** |
| AACGTTTGCTGCTGGAAACCATGCGTGGTATTCAGGCACTGTTTGTACGTCGCGACGAAGT GGAAGAAGCCTGGAAATGGGTAGACTCCATTACTGAGGCGTGGGCGATGGACAATGATG CGCCGAAACCGTATCAGGCCGGAACCTGGGGACCCGTTGCCTCGGTGGCGATGATTACCC GTGATGGTCGTTCCTGGAATGAGTTTGAGGGGGGTTCAGGCGGGTCGGGTGGCgtgagcaaggg cgaggagctgttcaccggggtggtgcccatcctggtcgagctggacggcgacgtaaacggccacaagttcagcgtgcgcggcgagggcgaggg cgatgccaccaacggcaagctgaccctgaagttcatctgcaccaccggcaagctgcccgtgccctggcccaccctcgtgaccaccctgacctacgg cgtgcagtgcttcagccgctaccccgaccacatgaagcgccacgacttcttcaagtccgccatgcccgaaggctacgtccaggagcgcaccatcag cttcaaggacgacggcacctacaagacccgcgccgaggtgaagttcgagggcgacaccctggtgaaccgcatcgagctgaagggcatcgacttc aaggaggacggcaacatcctggggcacaagctggagtacaacttcaacagccacaacgtctatatcaccgccgacaagcagaagaacggcatca aggccaacttcaagatccgccacaacgtggaggacggcagcgtgcagctcgccgaccactaccagcagaacacccccatcggcgacggccccg tgctgctgcccgacaaccactacctgagcacccagtccgtgctgagcaaagaccccaacgagaagcgcgatcacatggtcctgctggagttcgtga ccgccgccgggatcactcacggcatggacgagctgtacaagTAATGAATGATCGGCACGTAAGAGGTTCCAACTT TCACCATAATGAAATAAGATCACTACCGGGCGTATTTTTTGAGTTATCGAGATTTTCAGGA GCTAAGGAAGCTAAAATGGCCAAGCCTTTGTCTCAAGAAGAATCCACCCTCATTGAAAGA GCAACGGCTACAATCAACAGCATCCCCATCTCTGAAGACTACAGCGTCGCCAGCGCAGCT CTCTCTAGCGACGGCCGCATCTTCACTGGTGTCAATGTATATCATTTTACTGGGGGACCTT GTGCAGAACTCGTGGTGCTGGGCACTGCTGCTGCTGCGGCAGCTGGCAACCTGACTTGTAT CGTCGCGATCGGAAATGAGAACAGGGGCATCTTGAGCCCCTGCGGACGGTGCCGACAGGT GCTTCTCGATCTGCATCCTGGGATCAAAGCCATAGTGAAGGACAGTGATGGACAGCCGAC GGCAGTTGGGATTCGTGAATTGCTGCCCTCTGGTTATGTGTGGGAGGGCTAAGTAGGGAT AACAGGGTAATTATCTGCGCTTATCCTTTATGGTTATTTTACCGGTAACATGATCTTGCGCA GATTGTAGAACAATTTTTACACTTTCAGGCCTCGTGCGGATTCACCCACGAGGCTTTTTTT ATTACACTGACTGAAACGTTTTTGCCCTATGAGCTCCGGTTACAGGCGTTTCAGTCATAAA TCCTCTGAATGAAACGCGTTGTGAATC |
| **zwf-sfGFP-DAS+4-zeoR** |
| AACGTTTGCTGCTGGAAACCATGCGTGGTATTCAGGCACTGTTTGTACGTCGCGACGAAGT GGAAGAAGCCTGGAAATGGGTAGACTCCATTACTGAGGCGTGGGCGATGGACAATGATG CGCCGAAACCGTATCAGGCCGGAACCTGGGGACCCGTTGCCTCGGTGGCGATGATTACCC GTGATGGTCGTTCCTGGAATGAGTTTGAGGGGGGTTCAGGCGGGTCGGGTGGCgtgagcaaggg cgaggagctgttcaccggggtggtgcccatcctggtcgagctggacggcgacgtaaacggccacaagttcagcgtgcgcggcgagggcgaggg cgatgccaccaacggcaagctgaccctgaagttcatctgcaccaccggcaagctgcccgtgccctggcccaccctcgtgaccaccctgacctacgg cgtgcagtgcttcagccgctaccccgaccacatgaagcgccacgacttcttcaagtccgccatgcccgaaggctacgtccaggagcgcaccatcag cttcaaggacgacggcacctacaagacccgcgccgaggtgaagttcgagggcgacaccctggtgaaccgcatcgagctgaagggcatcgacttc aaggaggacggcaacatcctggggcacaagctggagtacaacttcaacagccacaacgtctatatcaccgccgacaagcagaagaacggcatca aggccaacttcaagatccgccacaacgtggaggacggcagcgtgcagctcgccgaccactaccagcagaacacccccatcggcgacggccccg tgctgctgcccgacaaccactacctgagcacccagtccgtgctgagcaaagaccccaacgagaagcgcgatcacatggtcctgctggagttcgtga ccgccgccgggatcactcacggcatggacgagctgtacaagGGTGGGGGTGGGAGCGGCGGCGGTGGCTCCGCG GCCAACGATGAAAACTATTCTGAAAACTATGCGGATGCGTCTTAATGAATGATCGGCACG TAAGAGGTTCCAACTTTCACCATAATGAAATAAGATCACTACCGGGCGTATTTTTTGAGTT ATCGAGATTTTCAGGAGCTAAGGAAGCTAAAATGGCCAAGCCTTTGTCTCAAGAAGAATC |

| |
|---|
| CACCCTCATTGAAAGAGCAACGGCTACAATCAACAGCATCCCCATCTCTGAAGACTACAG<br>CGTCGCCAGCGCAGCTCTCTCTAGCGACGGCCGCATCTTCACTGGTGTCAATGTATATCAT<br>TTTACTGGGGGACCTTGTGCAGAACTCGTGGTGCTGGGCACTGCTGCTGCTGCGGCAGCTG<br>GCAACCTGACTTGTATCGTCGCGATCGGAAATGAGAACAGGGGCATCTTGAGCCCCTGCG<br>GACGGTGCCGACAGGTGCTTCTCGATCTGCATCCTGGGATCAAAGCCATAGTGAAGGACA<br>GTGATGGACAGCCGACGGCAGTTGGGATTCGTGAATTGCTGCCCTCTGGTTATGTGTGGG<br>AGGGCTAAGTAGGGATAACAGGGTAATTATCTGCGCTTATCCTTTATGGTTATTTTACCGG<br>TAACATGATCTTGCGCAGATTGTAGAACAATTTTTACACTTTCAGGCCTCGTGCGGATTCA<br>CCCACGAGGCTTTTTTTATTACACTGACTGAAACGTTTTTGCCCTATGAGCTCCGGTTACA<br>GGCGTTTCAGTCATAAATCCTCTGAATGAAACGCGTTGTGAATC |
| **lpd-DAS+4-zeoR** |
| GCGGCGAGCTGCTGGGTGAAATCGGCCTGGCAATCGAAATGGGTTGTGATGCTGAAGACA<br>TCGCACTGACCATCCACGCGCACCCGACTCTGCACGAGTCTGTGGGCCTGGCGGCAGAAG<br>TGTTCGAAGGTAGCATTACCGACCTGCCGAACCCGAAAGCGAAGAAGAAGGCGGCCAAC<br>GATGAAAACTATTCTGAAAACTATGCGGATGCGTCTTAATAGCGAATCCATGTGGGAGTTT<br>ATTCTTGACACAGATATTTATGATATAATAACTGAGTAAGCTTAACATAAGGAGGAAAAA<br>CATATGTTACGCAGCAGCAACGATGTTACGCAGCAGGGCAGTCGCCCTAAAACAAAGTTA<br>GGTGGCTCAAGTATGGGCATCATTCGCACATGTAGGCTCGGCCCTGACCAAGTCAAATCC<br>ATGCGGGCTGCTCTTGATCTTTTCGGTCGTGAGTTCGGAGACGTAGCCACCTACTCCCAAC<br>ATCAGCCGGACTCCGATTACCTCGGGAACTTGCTCCGTAGTAAGACATTCATCGCGCTTGC<br>TGCCTTCGACCAAGAAGCGGTTGTTGGCGCTCTCGCGGCTTACGTTCTGCCCAAGTTTGAG<br>CAGCCGCGTAGTGAGATCTATATCTATGATCTCGCAGTCTCCGGCGAGCACCGGAGGCAG<br>GGCATTGCCACCGCGCTCATCAATCTCCTCAAGCATGAGGCCAACGCGCTTGGTGCTTATG<br>TGATCTACGTGCAAGCAGATTACGGTGACGATCCCGCAGTGGCTCTCTATACAAAGTTGG<br>GCATACGGGAAGAAGTGATGCACTTTGATATCGACCCAAGTACCGCCACCTAATTTTTCGT<br>TTGCCGGAACATCCGGCAATTAAAAAAGCGGCTAACCACGCCGCTTTTTTTACGTCTGCAA<br>TTTACCTTTCCAGTCTTCTTGCTCCACGTTCAGAGAGACGTTCGCATACTGCTGACCGTTGC<br>TCGTTATTCAGCCTGACAGTATGGTTACTGTC |
| **ydbKp-sfGFP** |
| TGATTGCAGTCCAGTTACGCTGGAGTCTGAGGCTCGTCCTGAATGATATCAAGCTTGAATT<br>CGTTCCTACAGCGTTTGCCGTTGGGTAATGCACACATCCCAATCGCCGTACCATCCAGTTG<br>ACGGGCAACAGAAAGCGAACCGCCGATCATTGCACAATTTGCTTCTCCACTACTGGACAT<br>CGACGCTTTTAAACCTGGCGCTACGTGCGCGGCAGTCGCCTGCTGAACAGGTTCACTACTA<br>CACGCCGACAACAATAAAGCGGCACACCCTACCCAAAACGCTGCTCGCATCTTTTCTTCCT<br>CTGATCTTCAAGCCAAACGACACCGCCATAAATAATAGGCAGCACAGAGGGCGGCGTCGA<br>GAGCTGTCCTGCGCGTTGCCCCGCATTTTTACTTTTTATGGCTATTTTTTTGCCCTCTGTTT<br>GATCAAAACATTCATTACGCTGATGTGGGGGACACAAAAGCGAAAATGCAGAAGAAAGC<br>CATTTGCTAAAATTGAAAGATTACTACTGGGCGCGCAGCAATTTCGTGCGCCCCTCATTCT<br>GTGTAGGAGGATAATCTATGGTGAGCAAGGGCGAGGAGCTGTTCACCGGGGTGGTGCCCA<br>TCCTGGTCGAGCTGGACGGCGACGTAAACGGCCACAAGTTCAGCGTGCGCGGCGAGGGCG<br>AGGGCGATGCCACCAACGGCAAGCTGACCCTGAAGTTCATCTGCACCACCGGCAAGCTGC<br>CCGTGCCCTGGCCCACCCTCGTGACCACCCTGACCTACGGCGTGCAGTGCTTCAGCCGCTA<br>CCCCGACCACATGAAGCGCCACGACTTCTTCAAGTCCGCCATGCCCGAAGGCTACGTCCA<br>GGAGCGCACCATCAGCTTCAAGGACGACGGCACCTACAAGACCCGCGCCGAGGTGAAGTT<br>CGAGGGCGACACCCTGGTGAACCGCATCGAGCTGAAGGGCATCGACTTCAAGGAGGACG<br>GCAACATCCTGGGGCACAAGCTGGAGTACAACTTCAACAGCCACAACGTCTATATCACCG<br>CCGACAAGCAGAAGAACGGCATCAAGGCCAACTTCAAGATCCGCCACAACGTGGAGGAC |

GGCAGCGTGCAGCTCGCCGACCACTACCAGCAGAACACCCCCATCGGCGACGGCCCCGTG
CTGCTGCCCGACAACCACTACCTGAGCACCCAGTCCGTGCTGAGCAAAGACCCCAACGAG
AAGCGCGATCACATGGTCCTGCTGGAGTTCGTGACCGCCGCCGGGATCACTCACGGCATG
GACGAGCTGTACAAGTAATGAGACGAATTCTCTAGATATCGCTCAATACTGACCATTTAA
ATCATACCTGACCTCC

### Section 2: Gene Silencing Arrays & Pathway Expression Constructs

#### a. pCASCADE Guide Array based Gene Silencing

The design and construction of CASCADE guides and guide arrays is illustrated below in Figure S7 and S8. The pCASCADE-control plasmid was prepared by swapping the pTet promoter in pcrRNA.Tet[15] with an insulated low phosphate induced ugpB promoter [4], as illustrated in Figure S7. Two promoters were responsible for regulating gltA gene, and sgRNA was designed for both promoters.[19] Four promoters were responsible for regulating gapA gene, and sgRNA was designed for the first promoter, since during exponential phase of growth, gapA mRNAs were mainly initiated at the highly efficient gapA P1 promoter and remained high during stationary phase compared to the other three gapA promoters.[20] Multiple promoters upstream of lpd gene were involved in lpd regulation (https://ecocyc.org/gene?orgid=ECOLI&id=EG10543#tab=showAll), thus design of unique and effective sgRNA for lpd only was not possible. Promoter sequences for fabI, udhA and zwf were obtained from EcoCyc database (https://ecocyc.org/). In order to design CASCADE guide array, CASCADE PAM sites near the -35 or -10 box of the promoter of interest were identified, 30 bp at the 3' end of PAM site was selected as the guide sequence and cloned into pCASCADE plasmid using Q5 site-directed mutagenesis (NEB, MA) following manufacturer's protocol, with the modification that 5% v/v DMSO was added to the Q5 PCR reaction. The pCASCADE-control vector was used as template. pCASCADE plasmids with arrays of two or more guides were prepared as described below and illustrated in Figure S8. The pCASCADE guide array plasmid was prepared by sequentially amplifying complementary halves of each smaller guide plasmid by PCR, followed by subsequent DNA assembly. Table S10 lists sgRNA guide sequences and primers used to construct them. All pCASCADE silencing plasmids are listed in Table S11 below and are available at Addgene.

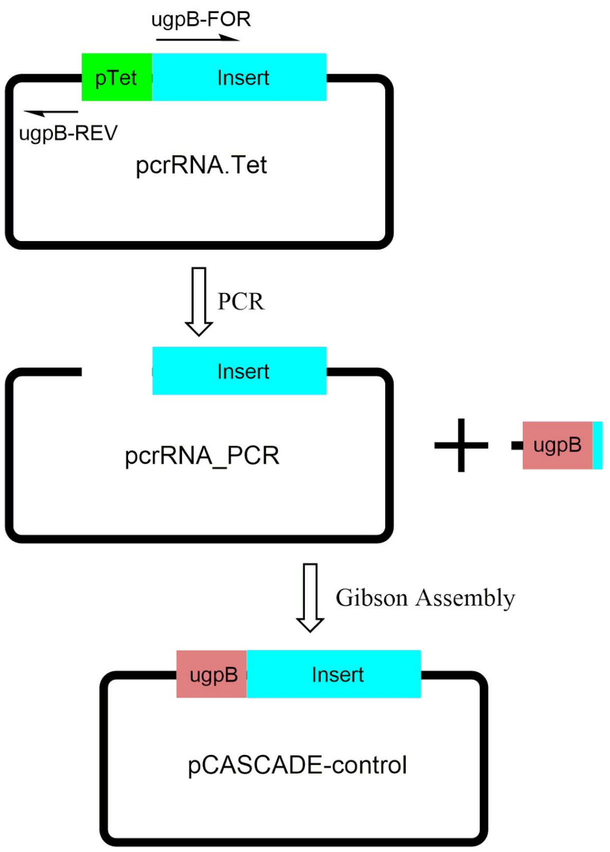


**Figure S7:** pCASCADE-control plasmid construction scheme.

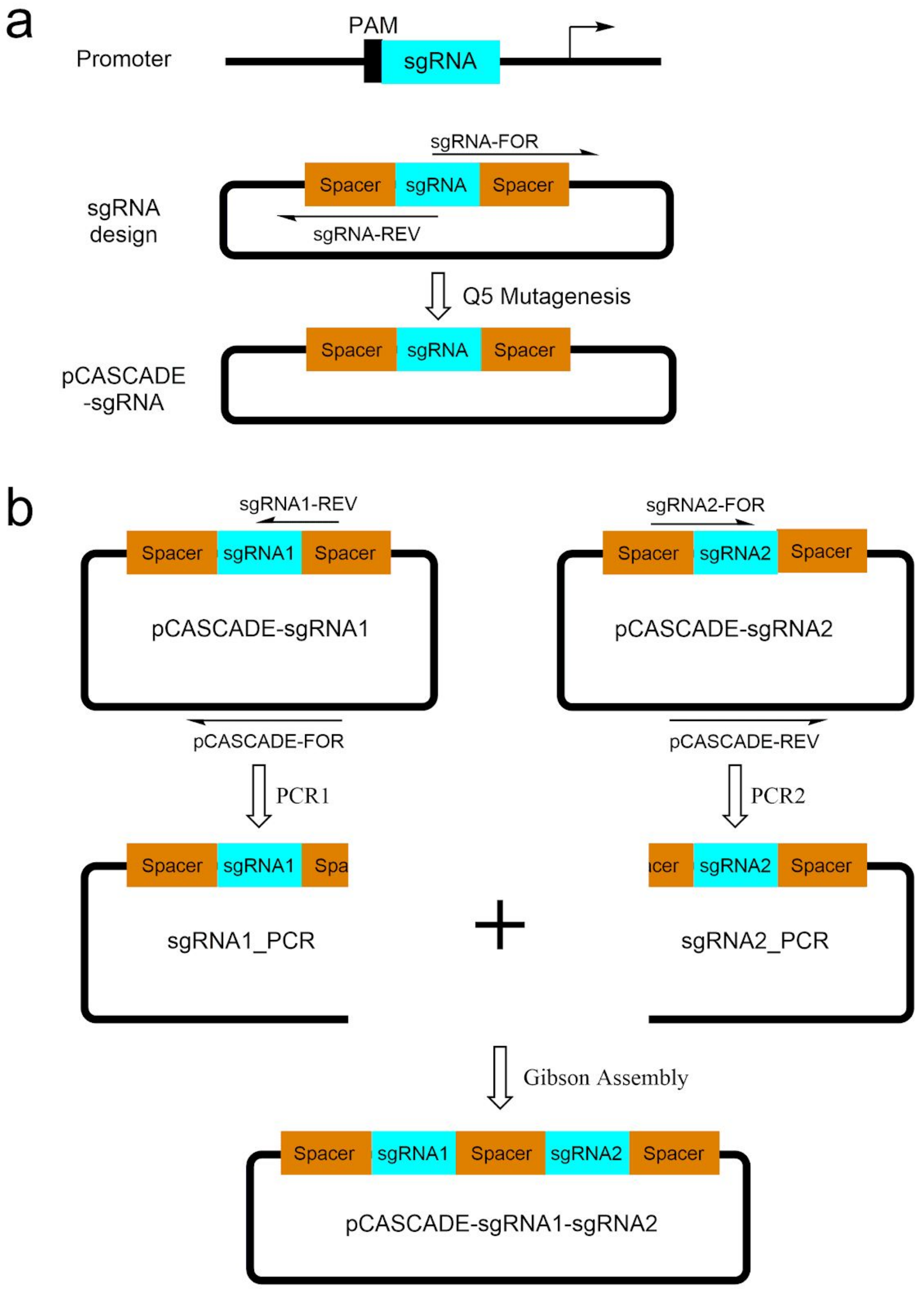


**Figure S8:** pCASCADE construction scheme. (a) single sgRNA cloning; (b) double sgRNA.

**Table S4:** List of sgRNA guide sequences and primers used to construct them. Spacers are italicized.

| sgRNA/Primer Name | Sequence | Template |
|---|---|---|
| gltA2 | *TCGAGTTCCCCGCGCCAGCGGGGATAAACCG*TATTGACCAATTCATTCGGGACAGTTATTAGT*TCGAGTTCCCCGCGCCAGCGGGGATAAACCG* | |
| gltA2-FOR | GGGACAGTTATTAGTTCGAGTTCCCCGCGCCAGCGGGGATAAACCGAAAAAAAAACCCC | pCASCADE ev |
| gltA2-REV | GAATGAATTGGTCAATACGGTTTATCCCCGCTGGCGCGGGGAACTCGAGGTGGTACCAGATCT | |
| proD | *TCGAGTTCCCCGCGCCAGCGGGGATAAACCG*AGTGGTTGCTGGATAACTTTACGGGCATGC*TCGAGTTCCCCGCGCCAGCGGGGATAAACCG* | |
| proD-FOR | AACTTTACGGGCATGCTCGAGTTCCCCGCGCCAGCGGGGATAAACCGAAAAAAAAACCCC | pCASCADE ev |
| proD-REV | ATCCAGCAACCACTCGGTTTATCCCCGCTGGCGCGGGGAACTCGAGGTGGTACCAGATCT | |
| zwf | *TCGAGTTCCCCGCGCCAGCGGGGATAAACCG*CTCGTAAAAGCAGTACAGTGCACCGTAAGA*TCGAGTTCCCCGCGCCAGCGGGGATAAACCG* | |
| zwf-FOR | CAGTGCACCGTAAGATCGAGTTCCCCGCGCCAGCGGGGATAAACCGAAAAAAAAACCCC | pCASCADE ev |
| zwf-REV | TACTGCTTTTACGAGCGGTTTATCCCCGCTGGCGCGGGGAACTCGAGGTGGTACCAGATC | |
| G2Z | *TCGAGTTCCCCGCGCCAGCGGGGATAAACCGTATTGACCAATTCATTC*GGGACAGTTATTAGT*TCGAGTTCCCCGCGCCAGCGGGGATAAACCG*CTCGTAAAAGCAGTACAGTGCACCGTAAGA*TCGAGTTCCCCGCGCCAGCGGGGATAAACCG* | |
| zwf-FOR | GCGCCAGCGGGGATAAACCGCTCGTAAAAG | pCASCADE-zwf |
| pCASCADE-REV | CTTGCCCGCCTGATGAATGCTCATCCGG | |
| pCASCADE-FOR | CCGGATGAGCATTCATCAGGCGGGCAAG | pCASCADE-G2 |
| gltA2-REV | CGGTTTATCCCCGCTGGCGCGGGGAACTCGAACTAATAACTGTC | |

**Table S5:** List of plasmids used in this study.

| Plasmid Utilized in this Study | | | | | |
|---|---|---|---|---|---|
| **Plasmid** | **Insert** | **Origin** | **Res** | **Addgene ID** | **Source** |
| pSIM5 | Recombineering genes | pSC101ts | Cm | NA | Court Lab |

| | | | | | |
|---|---|---|---|---|---|
| pSMART-HC-Kan | None - empty vector (ev) | ColE1 | Kan | NA | Lucigen |
| pcrRNA.Tet | gRNA control template | p15a | Cm | NA | Beisel Lab[2] |
| pCDF-ev | none control template | CloDF13 | Sp | 89596 | [1] |
| pSMART-GFPuv | yibDp-GFPuv | ColE1 | Kan | 65822 | [1] |
| pHCKan-yibDp-cimA3.7 | yibDp-cimA3.7 | ColE1 | Kan | 134595 | [1] |
| pHCKan-ydbKp-sfGFP | ydbKp-sfGFP | ColE1 | Kan | 159413 | this study |
| **Plasmid Constructed in this Study** | | | | | |
| **Plasmid** | **Insert** | **Origin** | **Res** | **Addgene ID** | **Source** |
| pCDF-mcherry1 | proDp-mCherry | CloDF13 | Sp | 87144 | this study |
| pCDF-mcherry2 | proDp-mCherry-DAS+4 | CloDF13 | Sp | 87145 | this study |
| pCASCADE-ev | empty gRNA control | p15a | Cm | 65821 | this study |
| pCASCADE-proD | proDp silencing gRNA | p15a | Cm | 65820 | this study |
| pCASCADE-G2 | gltA2p silencing gRNA | p15a | Cm | 65817 | this study |
| pCASCADE-Z | zwfp silencing gRNA | p15a | Cm | 65825 | this study |
| pCASCADE-G2Z | gltA2p, zwfp silencing gRNA array | p15a | Cm | 71338 | this study |

**Section 3: Dynamic Control over Protein Levels.**

Plasmids expressing fluorescent proteins and silencing guides were transformed into the corresponding hosts strain listed in Table S9. Strains were evaluated in triplicate in an m2p-labs Biolector™, which simultaneously measures fluorescence including GFPuv and mCherry levels, as well as biomass levels. Results are given in Figure 1.

**Table S6:** Strains used for Dynamic Control over protein levels

| RFP Strain | Plasmid | Host Strain |
|---|---|---|
| mCherry-control | pCDF-mcherry1 | DLF_Z002 |
| Proteolysis | pCDF-mcherry2 | DLF_Z0025 |
| Silencing | pCDF-mcherry1+pCASCADE-proD | DLF_Z01517 |
| Proteolysis+Silencing | pCDF-mcherry2+pCASCADE-proD | DLF_Z0025 |

OD600 readings were corrected using the formula below, where OD600 refers to an offline measurement, OD600* refers to Biolector biomass reading, $t_0$ indicates the start point, and $tf$ indicates the final point.

$$Equation\ S1:\ OD600_t = (OD600^*_t - OD600^*_{t0}) * \frac{(OD600_{tf} - OD600_{to})}{(OD600^*_{tf} - OD600^*_{to})} + 0.25$$

**Section 4: FGM 3 Media**

**Media Stock Solutions**

- 10X concentrated Ammonium-Citrate 30 salts (1L) by mixing 30 g of $(NH_4)_2SO_4$ and 1.5 g Citric Acid in water with stirring, adjust pH to 7.5 with NaOH. Autoclave and store at room temperature (RT).
- 10X concentrated Ammonium-Citrate 90 salts (1L) by mixing 90 g of $(NH_4)_2SO_4$ and 2.5 g Citric Acid in water with stirring, adjust pH to 7.5 with NaOH. Autoclave and store at RT.
- 1 M Potassium 3-(N-morpholino) propanesulfonic Acid (MOPS), adjust to pH 7.4 with KOH. Filter sterilize (0.2 μm) and store at RT.
- 0.5 M potassium phosphate buffer, pH 6.8 by mixing 248.5 mL of 1.0 M $K_2HPO_4$ and 251.5 mL of 1.0 M $KH_2PO_4$ and adjust to a final volume of 1000 mL with ultrapure water. Filter sterilize (0.2 μm) and store at RT.
- 2 M $MgSO_4$ and 10 mM $CaSO_4$ solutions. Filter sterilize (0.2 μm) and store at RT.
- 50 g/L solution of thiamine-HCl. Filter sterilize (0.2 μm) and store at 4°C.
- 500 g/L solution of glucose, dissolving by stirring with heat. Cool, filter sterilize (0.2 μm), and store at RT.
- 500X Trace Metal Stock: Prepare a solution of micronutrients in 1000 mL of water containing 10 mL of concentrated $H_2SO_4$. 0.6 g $CoSO_4 \cdot 7H_20$, 5.0 g $CuSO_4 \cdot 5H_20$, 0.6 g $ZnSO_4 \cdot 7H_20$, 0.2 g $Na_2MoO_4 \cdot 2H_2O$, 0.1 g $H_3BO_3$, and 0.3 g $MnSO_4 \cdot H_2O$. Filter sterilize (0.2 μm) and store at RT in the dark.
- Prepare a fresh solution of 40 mM ferric sulfate heptahydrate in water, filter sterilize (0.2 μm) before preparing media each time.

**Media Components:** Prepare the final working medium by aseptically mixing stock solutions based on the following tables in the order written to minimize precipitation, then filter sterilize (with a 0.2 μm filter).

**Table S7:** ***FGM3 Media, pH 6.8:***

| Ingredient | Concentration Stock | Volume in 1 L (mL) | Final Concentration |
|---|---|---|---|
| Ammonium-Citrate 30 Salts, pH 7.5 | 10 X | 100.0 | 1 X |
| Phosphate Buffer, pH 6.8 | 500 mM | 3.6 | 1.80 mM |
| Trace Metals | 500 X | 2.0 | 1 X |
| Fe (II) Sulfate | 40 mM | 2.0 | 0.08 mM |
| $MgSO_4$ | 2 M | 1.0 | 2.00 mM |
| $CaSO_4$ | 10 mM | 5.0 | 0.05 mM |
| Glucose | 500 g/L | 90.0 | 45 .0 g/L |

| | | | |
|---|---|---|---|
| MOPS | 1 M | 200.0 | 200 mM |
| Thiamine-HCl | 50 g/L | 0.2 | 0.01 g/L |

### Section 5: Additional Results

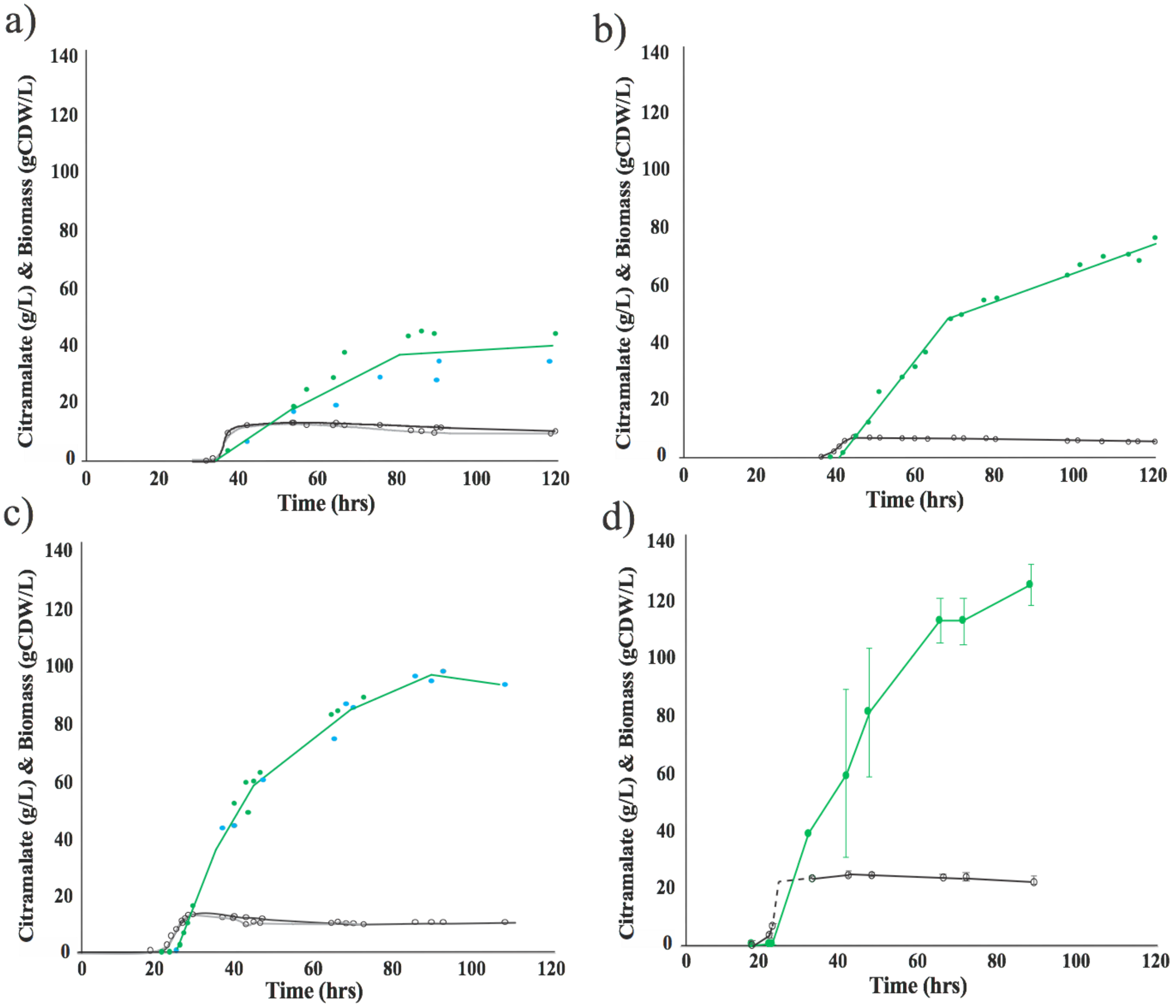


**Figure S9:** Citramalate and biomass production were measured for the control strain (a) and the "G" valve strain (b) and the "GZ" valve strain (c) in fermentations targeting biomass levels of 10gCDW/L. Duplicate runs, biomass levels in gray and black, citramalate titers in green and blue. (d) Citramalate production and biomass levels in fermentations targeting biomass levels of 25 gCDW. The average of triplicate runs, biomass black and citramalate green. Dashed line represents extrapolated growth due to missed samples.

### References


1. Menacho-Melgar, R. *et al.* Scalable, two-stage, autoinduction of recombinant protein expression in E. coli utilizing phosphate depletion. *Biotechnol. Bioeng.* **26**, 44 (2020).
2. Luo, M. L., Mullis, A. S., Leenay, R. T. & Beisel, C. L. Repurposing endogenous type I CRISPR-Cas systems for programmable gene repression. *Nucleic Acids Res.* **43**, 674–681 (2015).
